# Host phylogeny and ecological associations best explain *Wolbachia* host shifts in scale insects

**DOI:** 10.1101/2021.10.01.462721

**Authors:** Ehsan Sanaei, Gregory F Albery, Yun Kit Yeoh, Yen-Po Lin, Lyn G Cook, Jan Engelstädter

## Abstract

*Wolbachia* are among the most prevalent and widespread endosymbiotic bacteria on earth. *Wolbachia*’ s success in infecting an enormous number of arthropod species is attributed to two features: the range of phenotypes they induce in their hosts, and their ability to switch to new host species. Whilst much progress has been made in elucidating the phenotypes induced by *Wolbachia*, our understanding of *Wolbachia* host shifting is still very limited: we lack answers to even fundamental questions concerning *Wolbachia*’s routes of transfer and the importance of factors influencing host shifts. Here, we investigate the diversity and host-shift patterns of *Wolbachia* in scale insects, a group of arthropods with intimate associations with other insects that make them well-suited to studying host shifts. Using Illumina pooled amplicon sequencing of *Wolbachia*-infected scale insects and their direct associates we determined the identity of all *Wolbachia* strains, revealing that 32% of samples were multiply infected (with up to five distinct strains per species). We then fitted a Generalised Additive Mixed Model (GAMM) to our data to estimate the influence of factors such as the host phylogeny and the geographic distribution of each species on *Wolbachia* strain sharing among scale insect species. The model predicts no significant contribution of host geography but strong effects of host phylogeny, with high rates of *Wolbachia* sharing among closely related species and a sudden drop-off in sharing with increasing phylogenetic distance. We also detected the same *Wolbachia* strain in scale insects and several intimately associated species (ants, wasps, beetles, and flies). This indicates putative host shifts and potential routes of transfers via these associates and highlights the importance of ecological connectivity in *Wolbachia* host-shifting.

## Introduction

*Wolbachia* is one of the best-known groups of heritable endosymbionts on earth, widely distributed in arthropods and some nematodes (Hertig, 1936; Sironi et al., 1995; Werren, 1997). These bacteria form one of the most abundant and diverse groups of symbionts on earth: an estimated 40-60% of arthropod species are infected with *Wolbachia* strains (Zug & Hammerstein, 2012; Weinert et al., 2015). Their ability to induce various forms of reproduction manipulations (Rousset et al., 1992; Werren et al., 2008), and applications in such as the control of vector born disease (Kambris et al., 2009; Hoffmann et al., 2015; Ross et al., 2019), are key aspects of *Wolbachia* studied in the last two decades.

Like many other symbionts, the current distribution of *Wolbachia* results from three major processes: co-diversification with the host clade, shifting between host species, and symbiont loss (Thompson, 1987; Charleston & Perkins, 2006). Although co-speciation is common among *Wolbachia* strains belonging to supergroups C and D in their nematode hosts (Bandi et al., 1998; Fenn & Blaxter, 2004) and certain strains of supergroup F infecting bed bugs (Balvín et al., 2018)), many studies have failed to find evidence of codiversification between *Wolbachia* strains of supergroups A/B and arthropods (e.g., in fig wasps (Shoemaker et al., 2002), ants (Frost et al., 2010), butterflies (Ahmed et al., 2016), bees (Gerth et al., 2013), and collembolans (Ma et al., 2017)). In the absence of co-diversification, host-shifting is the alternative hypothesis to explain the current distribution of *Wolbachia*, as reviewed in (Sanaei et al., 2021a). *Wolbachia* shift hosts when a given strain infects a novel arthropod species, mostly through horizontal transfer (Boyle et al., 1993; Heath et al., 1999) and possibly occasionally through hybridisation (Turelli et al., 2018; Cooper et al., 2019). Host shifting is thought to play a key role in explaining the current high prevalence, distribution and diversification of these bacteria. The possibility of host-shift events in *Wolbachia* has been confirmed through numerous transinfection studies when a strain is artificially introduced to an uninfected species (reviewed in Hughes & Rasgon, 2014), and the existence of “super spreader strains” that infect host species which are phylogenetically distantly related (e.g., ST41 strain type in Lepidoptera (Ilinsky & Kosterin, 2017)). Physical transfer of *Wolbachia* from donor to recipient species is the first step of host-shifting, achieved via various “routes of transfer” and usually facilitated by a biological vector or a suitable environmental medium (Vavre et al., 2003; Riegler et al., 2004). Routes of transfer reported so far include prey-predator interactions (Le Clec’h et al., 2013), host-parasite interactions (Cook & Butcher, 1999; Vavre et al., 1999; Ahmed et al., 2015) and sharing a common food resource (Li et al., 2017). Detecting and understanding the routes of transfer in a given *Wolbachia*-host system can improve our knowledge of both the mechanism of *Wolbachia* host-shifting and host ecology.

Host phylogeny and ecological connectivity are thought to be the two main factors determining *Wolbachia* host shifting. As phylogenetically closely related species are similar in many respects, including their intercellular environment and immunology (Perlman & Jaenike, 2003), it is expected that a given symbiont will shift more easily between them than between distantly related species (Charleston & Robertson, 2002). This assumption, referred to as the “phylogenetic distance effect” (PDE) (Longdon et al., 2011; Engelstädter & Fortuna, 2019), may partly explain host shifts of *Wolbachia* across closely related species. In spite of limited case studies which indicated the presence of PDE in part of the host phylogeny (e.g., in fig wasps (Shoemaker et al., 2002), fungus growing ants (Frost et al., 2010), bees (Gerth et al., 2013) and Collembolans (Ma et al., 2017)), the influence of PDE on *Wolbachia* host shifting is not clear. Overlapping geographic distributions of host species is another possible explanatory factor. Sharing a common habitat and consequently potential ecological interactions may lead to several direct and indirect physical contacts between a given donor and recipient host and, therefore, also increase the probability of *Wolbachia* host shifting. Indeed, several case studies documented host-shift events between host species that share the same habitat, e.g., in a rice field community (Kittayapong et al., 2003) and a mushroom habitat (Stahlhut et al., 2010)).

Here, we use scale insects as a model system to gain a better understanding of *Wolbachia* host shifting. With more than 8200 described species and 24 families, the superfamily of scale insects (Coccoidea) are globally distributed (Gullan & Cook, 2007; García Morales et al., 2016). Like many other members of the suborder Sternorrhyncha, such as aphids, whiteflies and psyllids, scale insects exclusively feed on plants and some are considered serious agricultural pests (Kondo et al., 2008). Scale insects have documented ecological associations with a range of arthropods species. In particular, many are usually observed in close interactions with ants through trophallaxis (when the scale insect’s produced honeydew is consumed by ants) (Hölldobler et al., 1990; Buckley & Gullan, 1991; Gullan et al., 1993). Despite several similarities with other hemipterans, Sanaei et al., (2021b) found that most species are predicted to have low to intermediate *Wolbachia* prevalence, in contrast to a u-shaped distribution predicted for most other groups (Hilgenboecker et al., 2008). Also, a positive correlation between *Wolbachia* infection in scale insects and their associate ants indirectly points to a plausible route of transfer (Sanaei et al., 2021b). These preliminary results provide a broad view of the *Wolbachia* infection dynamic in scale insects and thus enable us to investigate *Wolbachia* strains diversity and consequently host-shifting in scale insects. These insects are a suitable model to study *Wolbachia* host shifting for several reasons. First, we have a large collection of scale insect samples, with known *Wolbachia* infection status, covering much of the species diversity in Australia (29 species and 4 families). In addition, we have samples of scale insect direct ecological associates (such as ants and wasps), which is essential to study routes of *Wolbachia* transfer. Finally, we had crucial data of their geographic distribution in Australia which can enable us to investigate the role of host geographic distance effect in *Wolbachia* horizontal transmission.

Studying of *Wolbachia* host-shifting requires using and developing new methodologies. To overcome technical problems associated with Sanger sequencing (see Discussion), we adopted Illumina pooled-amplicon sequencing techniques to determine the *Wolbachia* strains in scale insects and their associate species. Using this effective methodology, we revealed the strain diversity and composition (from single to multiple infections) in scale insects. Using phylogenetic trees of both scale insects and their *Wolbachia* strains, and the geographic distribution range of each scale insect species, we assessed which factors (phylogeny or geography) best explain host-shifting events. Finally, by determining *Wolbachia* strains in individual scale insects and directly associated individuals of other species, we identified plausible routes of horizontal transfer.

## Materials and Methods

### Sampling

We selected 59 specimens from 29 scale insect species that had tested positive for *Wolbachia* in a previous *Wolbachia* screening project (Sanaei et al., 2021b) (File S2). 16 of these scale insects were collected together with a directly associated insect (including ants, wasps, flies, beetles, moths), and these were also included. The tight ecological connection between the scale insect and an associate was established either by direct observation (e.g. ant-scale insect interactions) or by rearing both members of the pair in the laboratory conditions (e.g. rearing parasitoids from the scale insect sample). Based on observation, wasps and flies are mostly parasites and moth caterpillars are predators of scale insects. We selected 16 infected pairs: five scale insect-ant, seven scale insect-wasp, two scale insect-beetle, two scale insect-fly, and one scale insect-moth pair. We were unable to determine the species for any of the associates, except for the ants (*Technomyrmex albipes*) and one of the beetles (*Neopocadius pilistriatus*); however, we determined their COI barcode.

### PCR and sequencing

To retain all *Wolbachia* strains (especially in the case of co-infection) and consequently host-shifting events, we implemented an approach of Illumina pooled-amplicon sequencing. For this, we used 16S and MLST (Multilocus Sequence Typing) genes (Baldo et al., 2006), which included five housekeeping genes (coxA, fbpA, ftsZ, gatb, hcpA), as well as the *wsp (Wolbachia* Surface Protein) gene. Despite some limitation in using MLST (Bleidorn & Gerth, 2018), it is still a reliable source in strain determination and evolutionary history analysis (Wang et al., 2020). For the host genes, we targeted Cytochrome Oxidase I (COI), 18S and 28S Ribosomal RNA genes. The host genes were used later to confirm both scale insect and associate species identity and to build the host phylogeny. As a requirement for our Illumina sequencing platform, some of the primers were re-designed to yield products shorter than 500bp length (Table S1). We also added Illumina adaptors at the start (5’) of the forward and reverse primers (GTCTCGTGGGCTCGGAGATGTGTATAAGAGACAG and TCGTCGGCAGCGTCAGATGTGTATAAGAGACAG, respectively). PCR configurations were applied as originally suggested for each gene (Baldo et al., 2006).

We pooled the amplicons for all ten genes (seven *Wolbachia* and three host genes) per sample and sent them to the Australian Centre of Ecogenomics (The University of Queensland, Australia), where libraries were prepared using the Illumina AmpliSeq chemistry and sequenced on the Illumina MiSeq platform. The minimum total number of reads aimed for each sample was 10,000 and the read length for each direction was 300bp.

### Wolbachia strain determination

To determine the identity of the *Wolbachia* strain in our sample, we developed an R-based (R Core Team, 2013) bioinformatics pipeline based on the DADA2 pipeline (Callahan et al., 2016), which includes a series of quality controls, trimming and mapping to the references. In addition, we blasted all generated OTUs against the Genbank (https://www.ncbi.nlm.nih.gov/genbank/) and *Wolbachia* MLST database (https://pubmlst.org/organisms/Wolbachia-spp). The details of the pipeline are explained in File S1 (including Figure S1) and the R scripts are available in File S4.

### Reconstruction of phylogenies

*Wolbachia* and host genes were aligned in Geneious (Version 11.0.5, Biomatters) using the MAFFT algorithm (Katoh et al., 2002). Each gene was then trimmed to have identical lengths across samples. PartitionFinder2 (Lanfear et al., 2017) was used to find the best-fit partitioning scheme and substitution model for phylogeny estimation using default parameters. The results were then used as an input for estimating the Maximum Likelihood tree using RAxML (Stamatakis, 2014) with “Rapid Bootstrapping and Search for the Best scoring ML” and 1000 bootstrap replicates. As recombination is common among *Wolbachia* genes, the branch lengths of the *Wolbachia* phylogenetic tree were corrected with ClonalFrameML (Didelot & Wilson, 2015) to account for recombination events.

The MLST profile of all registered strains in the *Wolbachia* MLST depository (https://pubmlst.org/organisms/Wolbachia-spp) was downloaded (on 5^th^ November 2020). As most of the original *Wolbachia* MLST gene fragments were longer than the gene fragments in our study, the imported database was trimmed in Geneious to match the current study. The phylogenetic tree of all strains, including the reported strains in the MLST database and those from the current study, was estimated as above. This tree was used to determine the position of strains from scale insects within the various *Wolbachia* supergroups.

The phylogenetic trees of all hosts and *Wolbachia* strains, as well as the *Wolbachia*-host association network, were plotted in R by using the phytools package (Revell, 2012). A 3D interactive bipartite graph was also created using the bipartiteD3 R package (Terry, 2019). To test the phylogenetic congruence between *Wolbachia* and their hosts, we ran two tests. First, we performed a Parafit test (Legendre et al., 2002), which assesses the genetic distance similarity of host and parasite phylogenies. To this end, we used the parafit function implemented within the ape R-package (Paradis & Schliep, 2019) with the lingoes correction method for negative eigenvalues and 100,000 permutations. Second, we adopted the Procrustean Approach (known as PACo) which assesses the similarity between host and parasite trees by estimation of Euclidean embeddings derived from distance matrices (Balbuena et al., 2013). This test, which is implemented in PACo R package (Balbuena et al., 2013), was performed with 100,000 permutations. These two tests provide statistics to assess the independence of phylogenies by either rejecting or accepting the null hypothesis that the similarity between the trees is not higher than expected by chance. All R scripts developed and used in this study are provided in File S3.

### Factors determining host shifts

An expanded version of a Generalised Additive Mixed Model (GAMM), originally developed for viral sharing across mammal species (Albery et al., 2020), was applied by using the mgcv package in R (Wood, 2011). This GAMM allowed us to model a non-linear fit between our explanatory and response variables and allowed us to more readily account for their uneven distributions. Specifically, this model examined the probability of a given pair of scale insect species sharing one or more *Wolbachia* symbionts, as a function of their phylogenetic and geographic similarity, with a logistic link function:

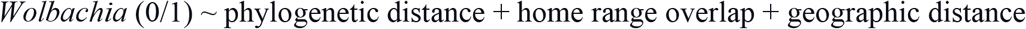

Phylogenetic distance was inferred from the Australian scale insect phylogenetic tree as explained above. To quantify habitat sharing between scale insect species, we constructed each species’ geographic range using their observed locations. For all species with 5 or more samples, we constructed a minimum convex polygon (MCP) in R. The coordinates for the MCP (File S4) were collected from various sources, mainly including the LGC collection (Cook Lab, School of Biological Science, The University of Queensland), ScaleNet (García Morales et al., 2016), the Atlas of Living Australia website (https://www.ala.org.au/), and several published articles (File S4). For each pair of species, we calculated the overlap of these polygons as a proportion of both species’ total range size. We also derived Euclidean distances between species’ sampling locations by calculating pairwise distances between species’ centroids. Species with fewer than 5 geographic observations were not included in the model. We also excluded *Coccus formicarii* which was collected from Taiwan and therefore difficult to fit in the model. A total number of 22 species were included in the GAMM model.

We fitted phylogenetic distance, home range overlap, and geographic distance as explanatory variables, and we fitted paired species’ identities as multi-membership random effects to account for variation in richness and sampling frequency between species (Albery et al., 2020). To quantify their impact on model fit we examined the change in deviance information criterion (DIC), where a change in 2 DIC was taken to represent an improved model. To avoid fitting too many variables in the model, we sequentially added each pairwise term, retaining the one that most improved model fit, and then repeating the process with the remaining variables, until no remaining variables improved the model. The R scripts are available in File S3.

## Results

The output of our bioinformatics pipeline based on Illumina pooled amplicon sequencing approach yielded read count numbers between 4658 and 36548 for *Wolbachia* genes per sample (for details see File S2). This high coverage enabled us to identify a diversity of *Wolbachia* strains within and across hosts. Among 75 *Wolbachia*-positive samples (59 scale insect samples and 16 associates), 68% were infected with a single *Wolbachia* strain and 32% were infected with more than one *Wolbachia* strain (20 double and 4 triple infected). Among 29 scale insect species that were screened in this study, 6 species were always found co-infected (File S2). Two screened samples of *Akermes scrobiculatus* were coinfected with strains belonging to Supergroup A (*w*Akel) and Supergroup B (*w*Ake2). The same pattern was observed for *Cystococcus echiniformis*, where two samples were coinfected with strains belonging to Supergroup A (*w*Sph1) and Supergroup F (*w*Cys1). A total of 63 strains were identified. Closely related strains (up to five bases difference across all seven genes) were grouped into “strain groups” (e.g., *w*Coc1). Given this, our determined strains clustered into 31 strain groups and belonged to three *Wolbachia*-supergroups (Figures 1, S2, S4). Most of the strains belong to Supergroups A (38) and B (21), but we also identified three strains from Supergroup F: *w*Cys1 and *w*Sph5 (respectively infecting *Cystococcus echiniformis* and *Sphaerococcus ferrugineus*), and *w*Sph3 (infecting two specimens of *S. ferrugineus*). Based on the MLST database, these are the first Supergroup F strains reported in Australia. Although *w*Cys1 is placed within Supergroup F, it forms a unique clade compared to other reported Supergroup F strains (Figure S2). The most diverse and abundant *Wolbachia* strain group in our dataset is *w*Sph1, which includes 12 closely related strains and was detected in 23 samples belonging to seven scale insect, four wasp and one ant species (Figure 2, S3 and S4, File S2). Similar strains which are grouped within *w*Sph1 were reported before in several Australian ant species (MLST ST = 54, 19, 478 and 112) (Russell, 2012). Based on the MLST database, it appears that this strain group has a cosmopolitan distribution (Oceania, North America, Europe, Asia and South Africa) and has already been reported in various insect groups (e.g. ST 19 in Coleoptera, Hymenoptera, Lepidoptera and Orthoptera). By contrast, some of the scale insect species are infected with unique *Wolbachia* strains that were not observed in other scale insects or reported in any other insects (by searching both MLST and GenBank on 5^th^ November 2020), including infection of *Apiomorpha variabilis* with *w*Aphi1 and co-infection of *Coccus hesperidium* with *w*Coc1 and *w*Coc2.

**Figure 1.**
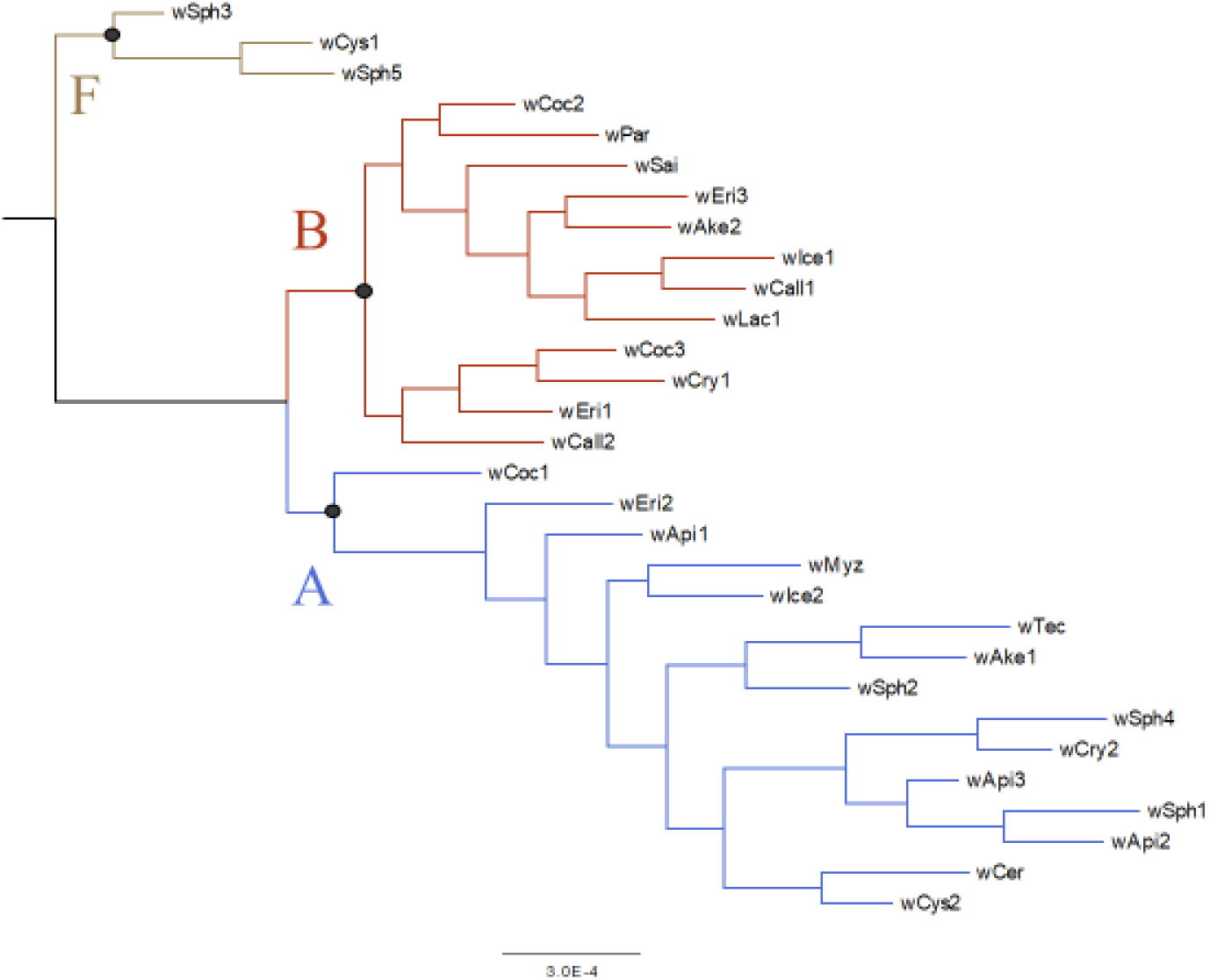
ML phylogenetic tree of the strain groups. Black circles are nodes with bootstrap value > 95%.

**Figure 2.**
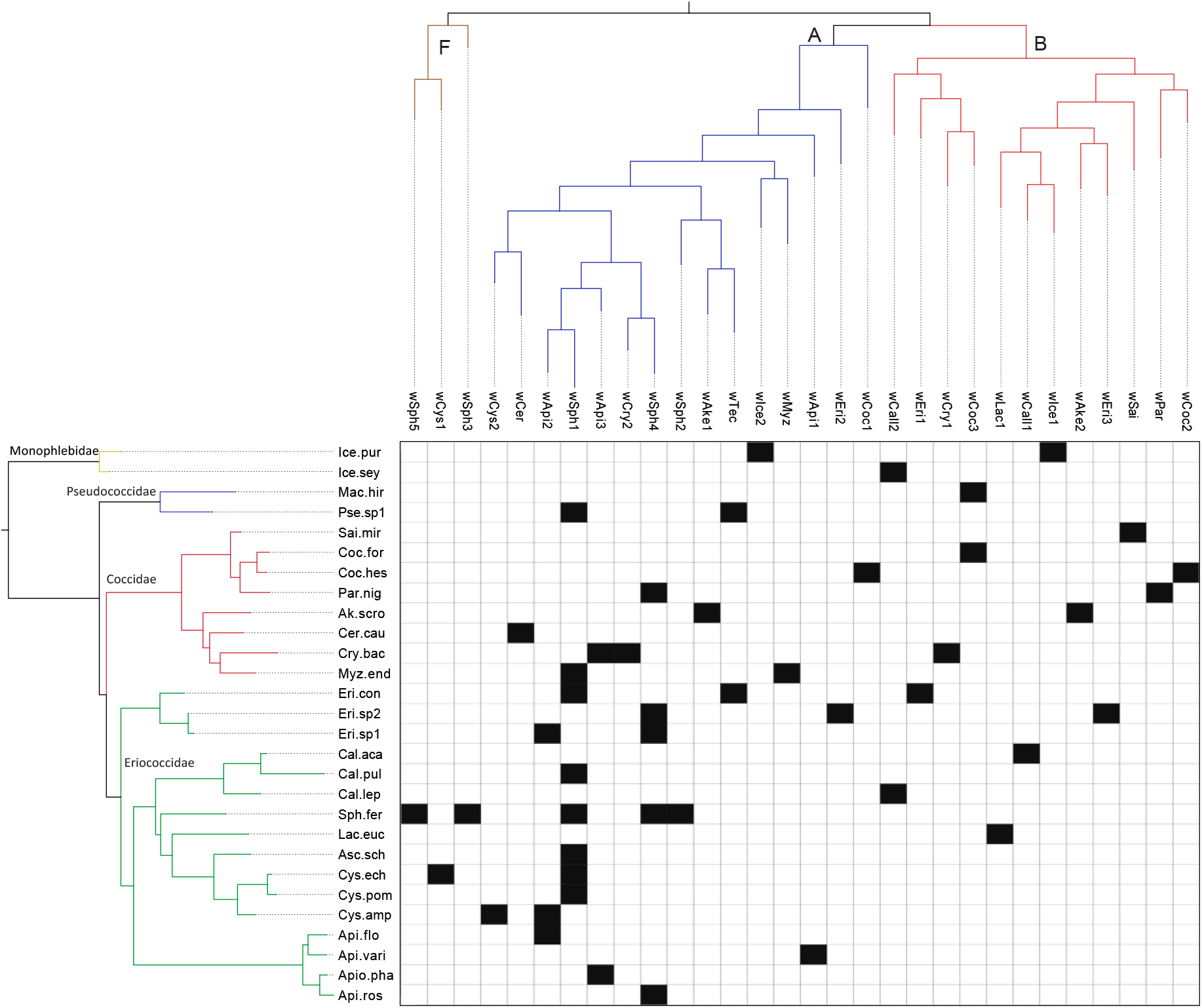
Phylogenetic tree of *Wolbachia* strain groups (top) and their scale insect hosts (left), with the black squares in the matrix indicating which host species are infected with which *Wolbachia* strain group(s). Colors in the phylogenies represent supergroups in *Wolbachia* and host families, respectively. Host species are represented by codes (for full species names please refer to File S2). An interactive tanglegram of the two phylogenetic trees is provided in File S5.

ML trees based on a 947 bp alignment of scale insect genes (including COI, 28S and 18S), and based on 2576 bp aligned *Wolbachia* genes (including MLST, wsp, and 16S) are shown in Figure 2. In addition, an interactive figure of *Wolbachia* sharing among all host species, including associates, is provided in File S5. Evidence of numerous host shifting events can be seen in these figures. Both the Parafit test (ParaFitGlobal = 0.0008, *P*-value = 0.27) and the PACo test (m^2^_XY_ = 49.3897, *P*-value = 0.2056) were non-significant. Therefore, there is no evidence for phylogenetic congruence between *Wolbachia* and their scale insect hosts.

Our *Wolbachia* sharing models revealed that incorporating phylogenetic distance substantially improved model fit (change in DIC = −5.88), and had a significant effect in the model (P<0.0001). The effect was highly non-linear, with high sharing probabilities at high relatedness that quickly dropped to near zero at greater phylogenetic distances (Figure 3). In contrast, incorporating geographic home range overlap slightly improved model fit (change in DIC = −2.76), but had no significant effect in the model (P=0.199). Inspecting the shape of the effect was not revealing. Furthermore, there was no significant effect of geographic distance between sampling locations (change in DIC > −2). Therefore, we do not interpret this effect as representing strong support for geographic effects on *Wolbachia* sharing.

**Figure 3.**
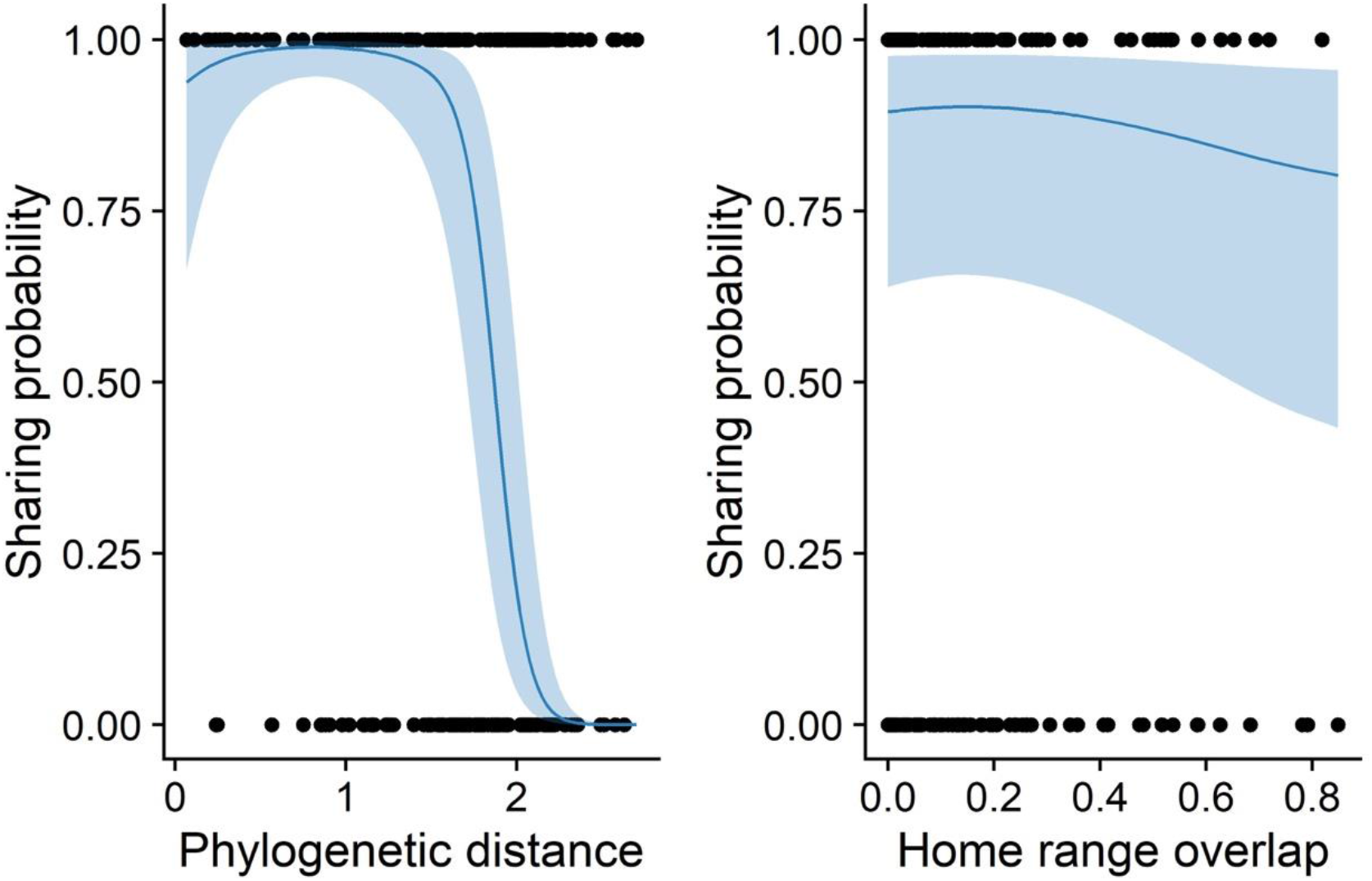
The effect of host phylogenetic distance (left) and home range overlap (right) on *Wolbachia* sharing probability. Points represent pairs of host species that either share (1) or do not share (0) the same *Wolbachia* strain; the thick blue line represents the mean predicted effect from our *Wolbachia* sharing GAMMs. The light blue ribbon represents the 95% confidence intervals of that effect. Sharing decreased with both phylogenetic and geographic distance, but the phylogenetic effect was significant and much steeper.

In searching for *Wolbachia* strains in pairs consisting of a scale insect and its direct associate, we found the same *Wolbachia* strain groups in one out of five ants, two out of two flies, one out of two beetles, and three out of seven wasps. We did not find the same strain group in the single tested moth-scale insect pair. As in scale insects overall, the most common strain group shared between scale insects and associates is the *w*Sph1 strain group (File S2).

## Discussion

### Amplicon sequencing as a powerful method of *Wolbachia* strain determination

Strain determination is a key step in studying *Wolbachia* distribution and host-shifting among a given host group that needs to be performed using an efficient method. Given that infection with more than one *Wolbachia* strain is common in various arthropod groups (Werren et al., 1995; Perrot-Minnot et al., 1996; Hiroki et al., 2004; Narita et al., 2007; Hou et al., 2020), strain determination methods should be able to distinguish and identify strains in both singly and multiply-infected samples. The traditional method of using Sanger sequencing is not effective in dealing with co-infected arthropod samples, and improvements such as using different primers and cloning (Schuster, 2008; Vo & Jedlicka, 2014)) are costlier and more labour-intensive, and also have limitations (Van Borm et al., 2003; Schuler et al., 2011). High-throughput whole genome sequencing (WGS) would seem to be the most accurate available methodology for strain identification, but this approach has its own difficulties (Bleidorn & Gerth, 2018). First, given that *Wolbachia* is not culturable, it is challenging to obtain genetic material enriched for *Wolbachia* relative to host DNA, possibly resulting in low sequencing depth. Second, even with high sequencing depth, assembling *Wolbachia* genomes can be difficult due to a high density of mobile elements (Wang et al., 2019). Finally, the still relatively high costs of WGS make this approach less applicable in large *Wolbachia* surveys. Due to these limitations, only 33 *Wolbachia* annotated whole genomes have been publicly available on GenBank so far (as of October 2021). To overcome these technical obstacles, we suggest Illumina pooled amplicon sequencing as a middle-ground, efficient and affordable method that can be applied to large surveys and is also capable of dealing with multiple infections. In particular, the five *Wolbachia* MLST genes along with *wsp* and 16S used in our study appear to be well suited to distinguish between strains, as has also been shown in a recent comparative study of available whole genomes of *Wolbachia* (Wang et al., 2020).

### Wolbachia diversity in scale insects

This study revealed that a substantial portion of tested scale insects are infected with more than one strain of *Wolbachia* (27% double and 5% triple infected). We also found *Wolbachia* multiple infections in associate species (including wasps and ants), indicating co-infection might be a common phenomenon in most of these insect groups. However, it is important to caution that detecting a given *Wolbachia* strain in a given host is not conclusive evidence of a stable infection, and laboratory assays should be conducted to ascertain *Wolbachia* maternal transmission and establishment within the host population (Chrostek et al., 2017; Sanaei et al., 2021a). Moreover, in the case of parasitoids and predators, a detected strain may derive from their undigested prey rather than the screened insect itself. Unfortunately, laboratory rearing of collected samples is not feasible for large *Wolbachia* surveys such as the current study. Therefore, any interpretation from this type of data should be treated with caution.

Based on the MLST database (as of 31st August 2021), 24 strains with complete MLST gene sequences had previously been reported from the Australian fauna (https://pubmlst.org/organisms/wolbachia-spp). Here, we report 63 new strains (belonging to 31 strain groups) for Australia, including the first three Supergroup F strains in Australasia. Apart from two strains (*w*Sph4.1 = ST 289, and *w*Cal = ST 357), none of the strains in the current study were 100% identical to any registered in the MLST database. As our sequenced regions were slightly (~5%) smaller than the original MLST amplicons, there is a possibility that these two identical strains to the MLST profiles were different in the remaining part of the gene fragments. We found *w*Sph1 to be the most common and widely distributed strain group in Australia (detected in seven scale insects, four wasps and one ant species). Based on the phylogenetic tree of all reported strains in the MLST database and the current study strains, there are six registered strain group within the *w*Sph1 strain group (Figure S2). These strains seem to be globally distributed across various insect orders. For example, one of the strains in this group, registered as ST=19, has been reported in 16 different host species belonging to four insect orders. This broad host range may be an indicator of an extraordinary host-shifting ability of *w*Sph1. Mostly based on the number of infected host species, several *Wolbachia* strains have been reported with a similar ability (e.g. HVR-2 in ants (Tolley et al., 2019), ST41 in Lepidoptera (Ilinsky & Kosterin, 2017), and *w*Hypera in weevils (Sanaei et al., 2019)). Among all of the superspreaders, *w*Ri is one of the best-studied *Wolbachia* strain groups that has rapidly (within 14,000 years) naturally infected five *Drosophila* species (Turelli et al., 2018). *w*Ri can also be easily introduced to mosquitoes by transinfection, corroborating this strain’s potential to infect new host species (Fraser et al., 2017). Compared to *w*Ri, it seems that *w*Sph1 has been reported in a higher number of host species that are taxonomically more diversified (belonging to various insect orders). Although *w*Ri group has an extensive genomic diversity (Ishmael et al., 2009; Turelli et al., 2018), low variation (including four strains) has been observed in its MLST profiles (https://pubmlst.org/organisms/wolbachia-spp). Four strains have been reported in *w*Ri group and only one strain (ST=17) has been reported in more than one species of *Drosophila* genus (based on MLST website as of 31st August 2021).

Therefore, *w*Sph1 might have a higher diversity than *w*Ri and may therefore have a potential to be artificially introduced to other insects for human applications (e.g., controlling vector born disease). However, transinfection studies are necessary to ascertain the host-shifting ability of *w*Sph1 in laboratory conditions.

### Phylogenetic distance effect can explain host-shifting

As is typical of *Wolbachia* infection in an arthropod family, we could not find a signal of congruence between *Wolbachia* and scale insect phylogeny. Instead, the current distribution of *Wolbachia* in scale insects was most likely shaped by host-shifting. Among many potential factors determining host shifts, it seems that host phylogeny and geographic distributions are two major players (Sanaei et al., 2021a). Combining data from 25 transinfection studies, Russell *et al*. (2009) showed that there is a positive correlation between host phylogenetic relatedness and success of the *Wolbachia* transinfection. In addition, by focusing only on a part of the host phylogenetic tree, several studies uncovered a pattern of host-shifting among closely related species (Haine et al., 2005; Guz et al., 2012; Turelli et al., 2018). On the other hand, the observation of identical *Wolbachia* strains in species that live in the same area indicates the role of geography in host-shifting (Kittayapong et al., 2003; Stahlhut et al., 2010; Morrow et al., 2014; Gupta et al., 2021). The relative contributions of the host phylogenetic and geographic distance effect on *Wolbachia* host shifts are poorly understood (Sanaei et al., 2021a). Here, we tried to evaluate these two factors in *Wolbachia* host shifting by providing a powerful statistical method. The results of our GAMM indicate that host shifts in scale insects can be mainly explained by the phylogenetic distance effect (host shifting is more feasible between closely related species compared to distantly related) (Figure 3). This result is in line with numerous examples of finding the same *Wolbachia* strain group in congeneric species (e.g. *wHypera1* in the genus *Hypera* (Coleoptera) (Sanaei et al., 2019), *w*Lev in the genus *Lutzomyia* (Diptera) (Vivero et al., 2017), ST19 in the genus *Bicyclus* (Lepidoptera) (Duplouy & Brattström, 2018)).

Horizontal transfer of parasites/symbionts among closely related species can generate a phylogenetic signal similar to host-parasite co-speciation. However, there is indirect evidence advocating *Wolbachia* sharing patterns in scale insects that can be explained best by recent host-shifting. In contrast to horizontal transmission which occurs rapidly, co-speciation happens in an evolutionary timeframe which allows *Wolbachia* genes to be mutated. By investigating *Wolbachia* infection in *Nasonia* species complex, it is estimated that *Wolbachia* MLST genes mutation rate is one-third of their host nuclear genes (from nine single copy nuclear regions) (Raychoudhury et al., 2009). Although this ratio can be slightly different among various host species and *Wolbachia* strains, it can be adopted as a tool to distinguish co-diversifications from recent host shifting. Given that the lowest pairwise distance between host species nuclear genes that we have in our dataset is 2%, in the case of *Wolbachia* cospeciation, at least 17 bp differences (out of 2608 bp) should be observed between two closely related strains. We defined host-shift events based on sharing either identical strains or identical strain groups (which includes strains with up to only 5bp differences across all *Wolbachia* amplicons) (Figure S4 and File S2). In addition, in 73% of determined host-shift events in scale insects, shared *Wolbachia* strains have identical wsp which is less conserved compared to other MLST. Therefore, sharing identical strains or strain groups among scale insect species strongly supports the hypothesis of recent host shifts rather than co-speciation events.

As a first step in the host shift process, *Wolbachia* need to physically reach the recipient host species, which requires direct or indirect species interactions. Therefore, it is expected that one should observe host shifting among species with an overlap in geographic distribution. However, our model indicates that the geographic home range of scale insect species has no significant contribution to *Wolbachia* sharing (Figure 3). This finding may firstly relate to the age of infection. Estimation of the *Wolbachia* infection age and consequently the intervals of host-shift events is controversial, with vastly different estimates across different case studies being reported (from a few thousand (Turelli et al., 2018; Cooper et al., 2019) to nine million years (Bailly-Bechet et al., 2017), see also Sanaei et al (2021a)). If the changes in the host geographic distribution occur faster than *Wolbachia* host shift events, the current geographic distribution may not be able to explain host-shift events (and thus, we would need to reconstruct the historical home range). In addition, the geographic distribution of a given species is not necessarily representative of the realised niche of that species, including ecological connectivity (Pulliam, 2000; Kearney, 2006; Peterson & Soberón, 2012). Therefore, two species may have the same geographic distribution but have no direct or even indirect physical interactions (e.g. via sharing foods or other resources). In that case, host ecological niche may be a better tool to explain *Wolbachia* host-shifting. However, ecological niches are technically harder to measure, especially when trying to account for ecological interactions.

### Role of scale insect associate species in *Wolbachia* host shifts

Another possible reason why host geographic distance effect has such a weak effect on *Wolbachia* sharing is host shifting via ecological vectors. Such vectors can carry on the infection, either temporarily or permanently (Sanaei et al., 2021a), and transmit it to a recipient species at a distant geographic location from the donor species. From prey-predator (Johanowicz & Hoy, 1996; Le Clec’h et al., 2013) to host-parasitoid (Vavre et al., 1999; Kageyama et al., 2010; Tzuri et al., 2020) and trophallaxis interactions (Ramalho & Moreau, 2020), there are several direct and indirect ecological pathways which can be routes of *Wolbachia* transfer. Detection of these pathways provides a better understanding of the dynamics and global distribution of *Wolbachia* infections (Sanaei et al., 2021a). Intimacy of direct physical interaction between ants and scale insects may provide a route of microbial exchange, as seen in other hemipteran groups (Pringle & Moreau, 2017; Ivens et al., 2018). In a previous study, a positive correlation between *Wolbachia* infection in scale insects and their associates indicated that ants may play a role in host-shifting (Sanaei et al., 2021b). Here, we found that only one out of five case of infected ant-scale insect pairs shares the same *Wolbachia* strain. While we do not have enough statistical power to test which route of transfer is the most common and in which directions these transfers take place, our data support the hypothesis that the associates tested in the current study play a role in host shifting. Although positive correlations were not previously observed between infection of scale insects and their associates (Sanaei et al., 2021b), here we observed sharing similar *Wolbachia* strains between pairs of scale insect and not only ants but also wasps, beetles and flies (File S3). In addition, infection by the super-spreader stain “*w*Sph1” of several species of scale insects, ants, and wasps is another source of evidence for a substantial contribution of associate species in *Wolbachia* host-shifting in scale insects.

This study provided the first insight into *Wolbachia* strain diversity in scale insects, revealed a high portion of co-infected samples and detected *w*Sph1 as one the most common strain of *Wolbachia* in scale insects. We also found that the host phylogenetic distance effect plays a critical role in host shifting in scale insects. In future studies, the methodology suggested by this study could be applied to a larger data set to detect the factors influencing host-shifting on the global perspective.

## Acknowledgements

We are grateful to Sylvain Charlat, Benjamin Normark and Anne Duplouy for reading and reviewing the manuscript. We thank Juan Antonio Balbuena and Xavier Didelot for their technical advice on, respectively, PACo and ClonalFrame analysis. We also thank John Dwyer and Andrew Letten for providing constructive feedbacks on the statistical models. Penelope Mills kindly allowed us to access her scale insect database which is much appreciated.

## References

Ahmed M.Z., Breinholt J.W. & Kawahara A.Y. 2016. Evidence for common horizontal transmission of Wolbachia among butterflies and moths. BMC Evol. Biol. 16: 118.

Ahmed M.Z., Li S.-J., Xue X., Yin X.-J., Ren S.-X., Jiggins F.M., Greeff J.M. & Qiu B.-L. 2015. The intracellular bacterium Wolbachia uses parasitoid wasps as phoretic vectors for efficient horizontal transmission. PLoS Pathog. 11: e1004672.

Albery G.F., Eskew E.A., Ross N. & Olival K.J. 2020. Predicting the global mammalian viral sharing network using phylogeography. Nat. Commun. 11: 2260.

Bailly-Bechet M., Martins-Simões P., Szöllősi G.J., Mialdea G., Sagot M.-F. & Charlat S. 2017. How Long Does Wolbachia Remain on Board?. Mol. Biol. Evol. 34: 1183–1193.

Balbuena J.A., Míguez-Lozano R. & Blasco-Costa I. 2013. PACo: A Novel Procrustes Application to Cophylogenetic Analysis. PLOS ONE. 8: e61048.

Baldo L., Hotopp J.C.D., Jolley K.A., Bordenstein S.R., Biber S.A., Choudhury R.R., Hayashi C., Maiden M.C., Tettelin H. & Werren J.H. 2006. Multilocus sequence typing system for the endosymbiont Wolbachia pipientis. Appl. Environ. Microbiol. 72:7098–7110.

Balvín O., Roth S., Talbot B. & Reinhardt K. 2018. Co-speciation in bedbug Wolbachia parallel the pattern in nematode hosts. Sci. Rep. 8: 8797.

Bandi C., Anderson T.J.C., Genchi C. & Blaxter M.L. 1998. Phylogeny of Wolbachia in filarial nematodes. Proc. R. Soc. Lond. B Biol. Sci. 265: 2407–2413.

Bleidorn C. & Gerth M. 2018. A critical re-evaluation of multilocus sequence typing (MLST) efforts in Wolbachia. FEMS Microbiol. Ecol. 94: fix163.

Boyle L., O’Neill S.L., Robertson H.M. & Karr T.L. 1993. Interspecific and Intraspecific Horizontal Transfer of Wolbachia in Drosophila. Science. 260: 1796–1799.

Buckley R. & Gullan P. 1991. More Aggressive Ant Species (Hymenoptera: Formicidae) Provide Better Protection for Soft Scales and Mealybugs (Homoptera: Coccidae, Pseudococcidae). Biotropica. 23: 282–286.

Callahan B.J., McMurdie P.J., Rosen M.J., Han A.W., Johnson A.J.A. & Holmes S.P. 2016. DADA2: High-resolution sample inference from Illumina amplicon data. Nat. Methods. 13: 581–583.

Charleston M.A. & Perkins S.L. 2006. Traversing the tangle: Algorithms and applications for cophylogenetic studies. J. Biomed. Inform. 39: 62–71.

Charleston M.A. & Robertson D.L. 2002. Preferential Host Switching by Primate Lentiviruses Can Account for Phylogenetic Similarity with the Primate Phylogeny. Syst. Biol. 51: 528–535.

Chrostek E., Pelz-Stelinski K., Hurst G.D.D. & Hughes G.L. 2017. Horizontal Transmission of Intracellular Insect Symbionts via Plants. Front. Microbiol. 8: 2237.

Cook J.M. & Butcher R.D.J. 1999. The transmission and effects of Wolbachia bacteria in parasitoids. Res. Popul. Ecol. 41: 15–28.

Cooper B.S., Vanderpool D., Conner W.R., Matute D.R. & Turelli M. 2019. Wolbachia Acquisition by Drosophila yakuba-Clade Hosts and Transfer of Incompatibility Loci Between Distantly Related Wolbachia. Genetics. 212: 1399–1419.

Didelot X. & Wilson D.J. 2015. ClonalFrameML: Efficient Inference of Recombination in Whole Bacterial Genomes. PLOS Comput. Biol. 11: e1004041.

Duplouy A. & Brattström O. 2018. Wolbachia in the Genus Bicyclus: a Forgotten Player. Microb. Ecol. 75: 255–263.

Engelstädter J. & Fortuna N.Z. 2019. The dynamics of preferential host switching: Host phylogeny as a key predictor of parasite distribution. Evolution. 73: 1330–1340.

Fenn K. & Blaxter M. 2004. Are filarial nematode Wolbachia obligate mutualist symbionts?. Trends Ecol. Evol. 19: 163–166.

Fraser J.E., Bruyne J.T.D., Iturbe-Ormaetxe I., Stepnell J., Burns R.L., Flores H.A. & O’Neill S.L. 2017. Novel Wolbachia-transinfected Aedes aegypti mosquitoes possess diverse fitness and vector competence phenotypes. PLOS Pathog. 13: e1006751.

Frost C.L., Fernandez-Marin H., Smith J.E. & Hughes W.O.H. 2010. Multiple gains and losses of Wolbachia symbionts across a tribe of fungus-growing ants. Mol. Ecol. 19: 4077–4085.

García Morales M., Denno B.D., Miller D.R., Miller G.L., Ben-Dov Y. & Hardy N.B. 2016. ScaleNet: a literature-based model of scale insect biology and systematics. Database. 2016:.

Gerth M., Röthe J. & Bleidorn C. 2013. Tracing horizontal Wolbachia movements among bees (Anthophila): a combined approach using multilocus sequence typing data and host phylogeny. Mol. Ecol. 22: 6149–6162.

Gullan P.J., Buckley R.C. & Ward P.S. 1993. Ant-Tended Scale Insects (Hemiptera: Coccidae: Myzolecanium) within Lowland Rain Forest Trees in Papua New Guinea. J. Trop. Ecol. 9: 81–91.

Gullan P.J. & Cook L.G. 2007. Phylogeny and higher classification of the scale insects (Hemiptera: Sternorrhyncha: Coccoidea). Zootaxa. 413–425.

Gupta M., Kaur R., Gupta A. & Raychoudhury R. 2021. Are ecological communities the seat of endosymbiont horizontal transfer and diversification? A case study with soil arthropod community. Authorea. 5.

Guz N., Kocak E., Akpinar A., Gurkan M.O. & Kilincer A.N. 2012. Wolbachia infection in Trissolcus species (Hymenoptera: Scelionidae). Eur. J. Entomol. 109: 169–174.

Haine E.R., Pickup N.J. & Cook J.M. 2005. Horizontal transmission of Wolbachia in a Drosophila community. Ecol. Entomol. 30: 464–472.

Heath B.D., Butcher R.D.J., Whitfield W.G.F. & Hubbard S.F. 1999. Horizontal transfer of Wolbachia between phylogenetically distant insect species by a naturally occurring mechanism. Curr. Biol. 9: 313–316.

Hertig M. 1936. The Rickettsia, Wolbachia pipientis and Associated Inclusions of the Mosquito, Culex pipiens. Parasitology. 28: 453–486.

Hilgenboecker K., Hammerstein P., Schlattmann P., Telschow A. & Werren J.H. 2008. How many species are infected with Wolbachia? - a statistical analysis of current data. FEMS Microbiol. Lett. 281: 215–220.

Hiroki M., Tagami Y., Miura K. & Kato Y. 2004. Multiple infection with Wolbachia inducing different reproductive manipulations in the butterfly Eurema hecabe. Proc. R. Soc. Lond. B Biol. Sci. 271: 1751–1755.

Hoffmann A.A., Ross P.A. & Rašić G. 2015. Wolbachia strains for disease control: ecological and evolutionary considerations. Evol. Appl. 8: 751–768.

Hölldobler B., Holldobler F.P. of B.B., Wilson E.O. & Wilson H.C. in E. and U.R.P.E.E.O. 1990. The Ants. Harvard University Press, 784 pp.

Hou H.-Q., Zhao G.-Z., Su C.-Y. & Zhu D.-H. 2020. Wolbachia prevalence patterns: horizontal transmission, recombination, and multiple infections in chestnut gall wasp-parasitoid communities. Entomol. Exp. Appl. 168:.

Hughes G.L. & Rasgon J.L. 2014. Transinfection: a method to investigate Wolbachia-host interactions and control arthropod-borne disease. Insect Mol. Biol. 23: 141–151.

Ilinsky Y. & Kosterin O.E. 2017. Molecular diversity of Wolbachia in Lepidoptera: Prevalent allelic content and high recombination of MLST genes. Mol. Phylogenet. Evol. 109: 164–179.

Ishmael N., Hotopp J.C.D., Ioannidis P., Biber S., Sakamoto J., Siozios S., Nene V., Werren J., Bourtzis K., Bordenstein S.R. & Tettelin H. 2009. Extensive genomic diversity of closely related Wolbachia strains. Microbiology. 155: 2211–2222.

Ivens A.B.F., Gadau A., Kiers E.T. & Kronauer D.J.C. 2018. Can social partnerships influence the microbiome? Insights from ant farmers and their trophobiont mutualists. Mol. Ecol. 27: 1898–1914.

Johanowicz D.L. & Hoy M.A. 1996. Wolbachia in a Predator–Prey System: 16S Ribosomal Dna Analysis of Two Phytoseiids (Acari: Phytoseiidae) and Their Prey (Acari: Tetranychidae). Ann. Entomol. Soc. Am. 89: 435–441.

Kageyama D., Narita S., Imamura T. & Miyanoshita A. 2010. Detection and identification of Wolbachia endosymbionts from laboratory stocks of stored-product insect pests and their parasitoids. J. Stored Prod. Res. 46: 13–19.

Kambris Z., Cook P.E., Phuc H.K. & Sinkins S.P. 2009. Immune Activation by Life-Shortening Wolbachia and Reduced Filarial Competence in Mosquitoes. Science. 326: 134–136.

Katoh K., Misawa K., Kuma K. & Miyata T. 2002. MAFFT: a novel method for rapid multiple sequence alignment based on fast Fourier transform. Nucleic Acids Res. 30: 3059–3066.

Kearney M. 2006. Habitat, environment and niche: what are we modelling?. Oikos. 115: 186–191.

Kittayapong P., Jamnongluk W., Thipaksorn A., Milne J.R. & Sindhusake C. 2003. Wolbachia infection complexity among insects in the tropical rice-field community. Mol. Ecol. 12: 1049–1060.

Kondo T., Gullan P.J. & Williams D.J. 2008. Coccidology. The study of scale insects (Hemiptera: Sternorrhyncha: Coccoidea). Corpoica Cienc. Tecnol. Agropecu. 9:.

Lanfear R., Frandsen P.B., Wright A.M., Senfeld T. & Calcott B. 2017. PartitionFinder 2: New Methods for Selecting Partitioned Models of Evolution for Molecular and Morphological Phylogenetic Analyses. Mol. Biol. Evol. 34: 772–773.

Le Clec’h W., Chevalier F.D., Genty L., Bertaux J., Bouchon D. & Sicard M. 2013. Cannibalism and predation as paths for horizontal passage of Wolbachia between terrestrial isopods. PloS One. 8: e60232.

Legendre P., Desdevises Y. & Bazin E. 2002. A Statistical Test for Host–Parasite Coevolution. Syst. Biol. 51: 217–234.

Li S.-J., Ahmed M.Z., Lv N., Shi P.-Q., Wang X.-M., Huang J.-L. & Qiu B.-L. 2017. Plantmediated horizontal transmission of Wolbachia between whiteflies. ISME J. 11: 1019–1028.

Longdon B., Hadfield J.D., Webster C.L., Obbard D.J. & Jiggins F.M. 2011. Host Phylogeny Determines Viral Persistence and Replication in Novel Hosts. PLOS Pathog. 7: e1002260.

Ma Y., Chen W.-J., Li Z.-H., Zhang F., Gao Y. & Luan Y.-X. 2017. Revisiting the phylogeny of Wolbachia in Collembola. Ecol. Evol. 7: 2009–2017.

Morrow J.L., Frommer M., Shearman D.C.A. & Riegler M. 2014. Tropical tephritid fruit fly community with high incidence of shared Wolbachia strains as platform for horizontal transmission of endosymbionts. Environ. Microbiol. 16: 3622–3637.

Narita S., Nomura M. & Kageyama D. 2007. Naturally occurring single and double infection with Wolbachia strains in the butterfly Eurema hecabe: transmission efficiencies and population density dynamics of each Wolbachia strain. FEMS Microbiol. Ecol. 61: 235–245.

Paradis E. & Schliep K. 2019. ape 5.0: an environment for modern phylogenetics and evolutionary analyses in R. Bioinformatics. 35: 526–528.

Perlman S.J. & Jaenike J. 2003. Infection Success in Novel Hosts: An Experimental and Phylogenetic Study of Drosophila-Parasitic Nematodes. Evolution. 57: 544–557.

Perrot-Minnot M.-J., Guo L.R. & Werren J.H. 1996. Single and Double Infections with Wolbachia in the Parasitic Wasp Nasonia vitripennis Effects on Compatibility. Genetics. 143: 961–972.

Peterson A. & Soberón J. 2012. Species Distribution Modeling and Ecological Niche Modeling: Getting the Concepts Right. Nat. E Conserv. 10: 1–6.

Pringle E.G. & Moreau C.S. 2017. Community analysis of microbial sharing and specialization in a Costa Rican ant-plant-hemipteran symbiosis. Proc R Soc B. 284: 20162770.

Pulliam H.R. 2000. On the relationship between niche and distribution. Ecol. Lett. 3: 349–361.

R Core Team 2013. R: A language and environment for statistical computing.

Ramalho M.O. & Moreau C.S. 2020. The Evolution and Biogeography of Wolbachia in Ants (Hymenoptera: Formicidae). Diversity. 12: 426.

Raychoudhury R., Baldo L., Oliveira D.C. & Werren J.H. 2009. Modes of acquisition of Wolbachia: horizontal transfer, hybrid introgression, and codivergence in the Nasonia species complex. Evolution. 63: 165–183.

Revell L.J. 2012. phytools: an R package for phylogenetic comparative biology (and other things). Methods Ecol. Evol. 3: 217–223.

Riegler M., Charlat S., Stauffer C. & Merçot H. 2004. Wolbachia Transfer from Rhagoletis cerasi to Drosophila simulans: Investigating the Outcomes of Host-Symbiont Coevolution. Appl Env. Microbiol. 70: 273–279.

Ross P.A., Turelli M. & Hoffmann A.A. 2019. Evolutionary Ecology of Wolbachia Releases for Disease Control. Annu. Rev. Genet. 53: 93–116.

Rousset F., Bouchon D., Pintureau B., Juchault P. & Solignac M. 1992. Wolbachia endosymbionts responsible for various alterations of sexuality in arthropods. Proc R Soc Lond B. 250: 91–98.

Russell J.A. 2012. The ants (Hymenoptera: Formicidae) are unique and enigmatic hosts of prevalent Wolbachia (Alphaproteobacteria) symbionts. Myrmecol. News. 16: 18.

Russell J.A., Goldman-Huertas B., Moreau C.S., Baldo L., Stahlhut J.K., Werren J.H. & Pierce N.E. 2009. Specialization and geographic isolation among wolbachia symbionts from ants and lycaenid butterflies. Evolution. 63: 624–640.

Sanaei E., Charlat S. & Engelstädter J. 2021a. Wolbachia host shifts: routes, mechanisms, constraints and evolutionary consequences. Biol. Rev. 96: 433–453.

Sanaei E., Husemann M., Seiedy M., Rethwisch M., Tuda M., Toshova T.B., Kim M.J., Atanasova D. & Kim I. 2019. Global genetic diversity, lineage distribution, and Wolbachia infection of the alfalfa weevil Hypera postica (Coleoptera: Curculionidae). Ecol. Evol. 9: 9546–9563.

Sanaei E., Lin Y., Cook L.G. & Engelstaedter J. 2021b. Wolbachia in scale insects: a distinct pattern of infection frequencies and potential transfer routes via ant associates | bioRxiv. bioRxiv..

Schuler H., Arthofer W., Riegler M., Bertheau C., Krumböck S., Köppler K., Vogt H., Teixeira L.A.F. & Stauffer C. 2011. Multiple Wolbachia infections in Rhagoletis pomonella. Entomol. Exp. Appl. 139: 138–144.

Schuster S.C. 2008. Next-generation sequencing transforms today’s biology. Nat. Methods. 5: 16–18.

Shoemaker D.D., Machado C.A., Molbo D., Werren J.H., Windsor D.M. & Herre E.A. 2002. The distribution of Wolbachia in fig wasps: correlations with host phylogeny, ecology and population structure. Proc. R. Soc. Lond. B Biol. Sci. 269: 2257–2267.

Sironi M., Bandi C., Sacchi L., Di B.S., Damiani G. & Genchi C. 1995. Molecular evidence for a close relative of the arthropod endosymbiont Wolbachia in a filarial worm. Mol. Biochem. Parasitol. 74: 223–227.

Stahlhut J.K., Desjardins C.A., Clark M.E., Baldo L., Russell J.A., Werren J.H. & Jaenike J. 2010. The mushroom habitat as an ecological arena for global exchange of Wolbachia. Mol. Ecol. 19: 1940–1952.

Stamatakis A. 2014. RAxML version 8: a tool for phylogenetic analysis and post-analysis of large phylogenies. Bioinformatics. 30: 1312–1313.

Terry C. 2019. bipartiteD3: Interactive bipartite graphs.

Thompson J.N. 1987. Symbiont-induced speciation. Biol. J. Linn. Soc. 32: 385–393.

Tolley S.J.A., Nonacs P. & Sapountzis P. 2019. Wolbachia Horizontal Transmission Events in Ants: What Do We Know and What Can We Learn?. Front. Microbiol. 10: 296.

Turelli M., Cooper B.S., Richardson K.M., Ginsberg P.S., Peckenpaugh B., Antelope C.X., Kim K.J., May M.R., Abrieux A., Wilson D.A., Bronski M.J., Moore B.R., Gao J.-J., Eisen M.B., Chiu J.C., Conner W.R. & Hoffmann A.A. 2018. Rapid Global Spread of wRi-like Wolbachia across Multiple Drosophila. Curr. Biol. 28: 963–971.e8.

Tzuri N., Caspi-Fluger A., Betelman K., Rohkin Shalom S. & Chiel E. 2020. Horizontal Transmission of Microbial Symbionts Within a Guild of Fly Parasitoids. Microb. Ecol..

Van Borm S., Wenseleers T., Billen J. & Boomsma J.J. 2003. Cloning and sequencing of wsp encoding gene fragments reveals a diversity of co-infecting Wolbachia strains in Acromyrmex leafcutter ants. Mol. Phylogenet. Evol. 26: 102–109.

Vavre F., Fleury F., Lepetit D., Fouillet P. & Boulétreau M. 1999. Phylogenetic evidence for horizontal transmission of Wolbachia in host-parasitoid associations. Mol. Biol. Evol. 16:1711–1723.

Vavre F., Fouillet P. & Leury F. 2003. Between- and Within-Host Species Selection on Cytoplasmic Incompatibility-Inducing Wolbachia in Haplodiploids. Evolution. 57: 421–427.

Vivero R.J., Cadavid-Restrepo G., Herrera C.X.M. & Soto S.I.U. 2017. Molecular detection and identification of Wolbachia in three species of the genus Lutzomyia on the Colombian Caribbean coast. Parasit. Vectors. 10: 110.

Vo A.-T.E. & Jedlicka J.A. 2014. Protocols for metagenomic DNA extraction and Illumina amplicon library preparation for faecal and swab samples. Mol. Ecol. Resour. 14: 1183–1197.

Wang X., Xiong X., Cao W., Zhang C., Werren J.H. & Wang X. 2019. Genome Assembly of the A-Group Wolbachia in Nasonia oneida Using Linked-Reads Technology. Genome Biol. Evol. 11: 3008–3013.

Wang X., Xiong X., Cao W., Zhang C., Werren J.H. & Wang X. 2020. Phylogenomic Analysis of Wolbachia Strains Reveals Patterns of Genome Evolution and Recombination. Genome Biol. Evol. 12: 2508–2520.

Weinert L.A., Araujo-Jnr E.V., Ahmed M.Z. & Welch J.J. 2015. The incidence of bacterial endosymbionts in terrestrial arthropods. Proc. R. Soc. B Biol. Sci. 282: 20150249.

Werren J.H. 1997. Biology of wolbachia. Annu. Rev. Entomol. 42: 587–609.

Werren J.H., Baldo L. & Clark M.E. 2008. Wolbachia: master manipulators of invertebrate biology. Nat. Rev. Microbiol. 6: 741.

Werren J.H., Windsor D. & Guo L.R. 1995. Distribution of Wolbachia among neotropical arthropods. Proc. R. Soc. Lond. B Biol. Sci. 262: 197–204.

Wood S.N. 2011. Fast stable restricted maximum likelihood and marginal likelihood estimation of semiparametric generalized linear models. J. R. Stat. Soc. Ser. B Stat. Methodol. 73: 3–36.

Zug R. & Hammerstein P. 2012. Still a host of hosts for Wolbachia: analysis of recent data suggests that 40% of terrestrial arthropod species are infected. PloS One. 7: e38544.

