## Supplementary material for "Host phylogeny and ecological associations best explain *Wolbachia* host shifts in scale insects": File S1

### Supplementary tables, figures and text

Table S1. List of the primers used in the current study. For sequencing on the Illumina platform, we added Illumina adaptors at the start (5`) of the forward and reverse primers that are not shown in the table (GTCTCGTGGGCTCGGAGATGTGTATAAGAGACAG and TCGTCGGCAGCGTCAGATGTGTATAAGAGACAG, respectively).

| Locus | Size | Primer name | Sequences (5`-3`) F - R | References |
|---|---|---|---|---|
| 16S | 457 | Wol16SF3<br>Wol16SR3 | GCRAAGGCGTCTATCTGGT<br>TTCCTCCAGYTTATCACT | Sanaei *et al.* (2021) |
| gatB | 471 | gatB_F1<br>gatB_R1 | GAKTTAAAYCGYGCAGGBGTT<br>TGGYAAYTCRGGYAAAGATGA | Baldo *et al.* (2006) |
| coxA | 487 | coxA_F1<br>coxA_R1 | TTGGRGCRATYAACTTTATAG<br>CTAAAGACTTTKACRCCAGT | Baldo *et al.* (2006) |
| hcpA | 422 | IAhcpAF1<br>IAhcpNewR1 | CGYCTTCGCTCTGCTAT<br>GTTCTGGTTCTCCRAATTTTG | This study |
| ftsZ | 371 | IAftsZF2<br>IAftsZR1 | AGCAGGAATGGGCGGTGGTACT<br>GGATTRGATATTGCAGC | This study |
| fbpA | 492 | fbpAF<br>FbpAR | GAAATARCAGTTGCTGCAAA<br>GAAAGTYRAGCAAGYTCTG | Baldo *et al.* (2006) |
| wsp | ~365 | IAwspF2<br>IAwspR1 | GCATTTGGTTAYAAAATGG<br>ACATCATARCTAACACCAGC | This study |
| COI | 365 | COI_mlCOIintF<br>COI_Fol-degen-rev2 | GGWACWGGWTGAACWGTWTAYCCYCC<br>TANACYTCNGGRTGNCCRAARAAYCA | Krehenwinkel *et al.* (2018) |
| 18S | ~300 | 18S_2880F<br>18S_E10R | CTGGTTGATCCTGCCAGTAG<br>CGGTTTTGATCTAATAAGAGC | von Dohlen & Moran (1995) |
| 28S | ~300 | 28S_D2F<br>28S_D2R | AGAGAGAGTTCAAGAGTACGTG<br>TTGGTCCGTGTTTCAAGACGGG | Gimeno *et al.* (1997) |

##### Generating OTUs

All raw sequences in fastq format were registered and deposited in the GenBank as biosamples, and their accession numbers (SAMN21874044 to SAMN21874118) are available in File S2. To determine *Wolbachia* strains, we developed an R-based bioinformatic procedure from the DADA2 pipeline (Callahan et al., 2016). After extraction and correction of sample names, quality plots were generated for forward and reverse reads using FastQC (Brown et al., 2017). In almost all samples there was a low-quality region (Phred score < 25) at the start of forward and the end of reverse sequences. Based on this pattern, we applied a general trimming rule to trim 20bp from the start and 30bp from the end of all reads by trimmomatic (Bolger et al.,

2014). Then, we merged forward and reverse reads by BBmerge (Bushnell et al., 2017). After removing chimeras by consensus method (Callahan et al., 2016), we generated the OTU (Operational Taxonomic Unit) table including unique sequences and their coverage (counts). To assign the gene for each unique variant, OTUs are mapped to the *Wolbachia* reference genes and host-genes (provided from the pilot Sanger sequencing of each gene separately (Sanaei et al., 2021). The remaining reads, including those *Wolbachia* genes that did not hit to the references or the host genes (18S, 28S and COI) were blasted against the NCBI and *Wolbachia* MLST database. The remaining unknown OTUs that failed to match with any relevant arthropod or bacteria genes were discarded. In very few cases (12 amplicons out of 225), sequences of some of the scale insect genes were not obtained from the output of Illumina sequencing (Red cells in the "strains for R input" tab in File S2). In that case, we adopted those missed genes from sequences of that species (from Australia) available in GenBank to re-construct the phylogeny (File S2). Also, in the case of missing genes from a *Wolbachia* strain in a given sample, we adopted a replacement from the same strain (strain group) on other samples or, in two cases, from the closest MLST profile (File S2). All R scripts are provided in File S3.

**Strain determination**

In the case of a single infection (existence of a single unique sequence variant per *Wolbachia* gene), *Wolbachia* strain determination was conducted by assigning each mapped (or blasted) variant to its corresponding *Wolbachia* gene. Due to duplication, recombination, PCR and sequencing errors, it is common to observe more than one OTU per gene (even in single infected samples). Moreover, it is highly possible to observe OTUs with length variation but identical overlap regions. Therefore, strain determination was manually performed by respectively following the criteria below (a summary is provided in Figure S1):

1. A sample is assigned "single infected" if a maximum of three out of seven *Wolbachia* genes have multiple distinct OTUs.

2. In a given gene of a single strain infected sample, OTUs with less than 5% total coverage of all OTUs of that gene, were considered sequencing errors and removed.

3. In the case of having closely related OTUs with equal lengths in a single infected sample: OTUs with a maximum of two bases differences assigned to a given gene were merged to form

a consensus OTU. This procedure was commonly performed for the wsp gene when usually more than one OTU per gene is observed.

4. In the case of having closely related or identical OTUs with unequal lengths in a single infected sample: If the identical overlap region is more than 80% of the total length of all similar OTUs and the less abundant variant have more than 5% of the total coverage for that gene, the OTUs are be merged to create a new OTU. As a result, the merged sequence would have a slightly larger length than typical sequences. However, to define the strain and build the phylogenetic tree, the extra length would be trimmed. This procedure was conducted for the coxA gene when length variation was common.

5. In the case of multiple infections (double or triple): By mapping to the strain references, OTUs from the double or triple infected samples can be distinguished and separated. At first, the strains of the single infected samples were used as references. In the absence of any matches, the OTUs were blasted against the *Wolbachia* MLST database. In the case of matching OTUs to the reported strains (identical or maximum three bp differences), that reported strain is applied as a reference strain. However, 16S is not part of the MLST protocol and rarely reported for strain-description purposes. Therefore, the phylogenetic tree of 16S OTUs was reconstructed and compared to the MLST phylogenetic trees to assign 16S to the correct strain. In the absence of any references, if the sets of *Wolbachia* genes belong to different supergroups (A vs. B) or are located very far from each other in the phylogenetic tree, the strain determination becomes feasible. If the mentioned criteria failed, the unsorted remaining *Wolbachia* strains in a co-infected sample are being reported.

Assigned *Wolbachia* strains were named from the first three letters of the genus name of the first host from which they were diagnosed.

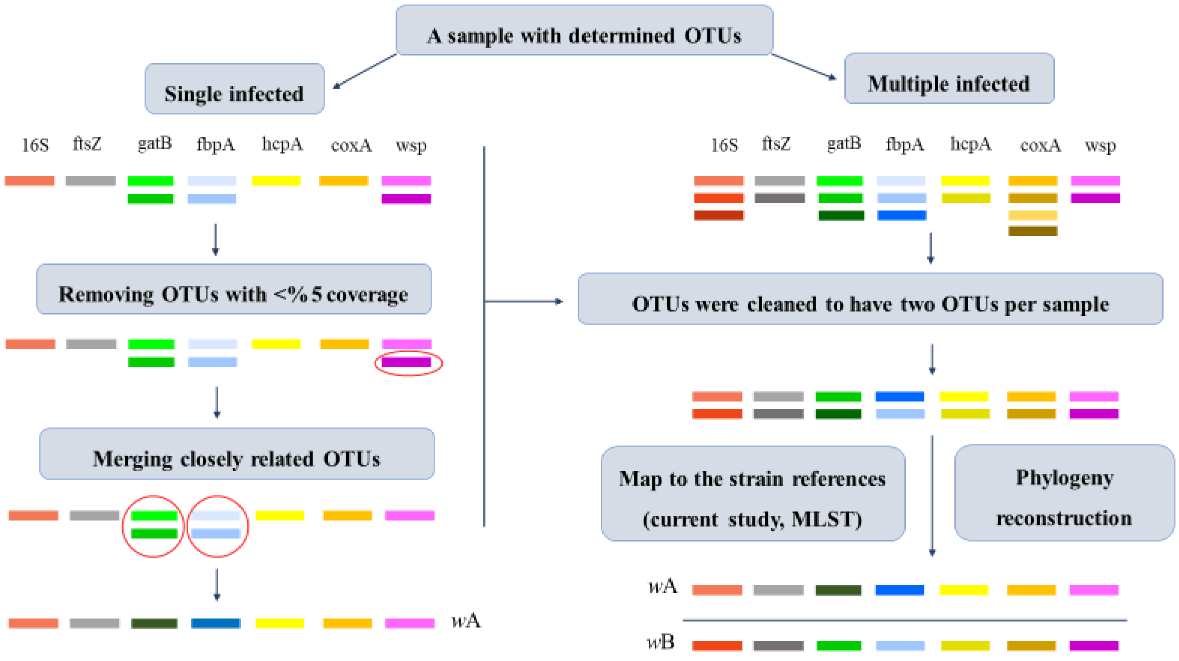


Figure S1. Summary of strain determination processes from generated OTUs in single and multiple (here an example of double) infected samples. OTUs in each *Wolbachia* gene are distinguishable by their colour. By following these steps, a given strain *w*A in single infected and *w*A/*w*B in double infected sample can be determined.

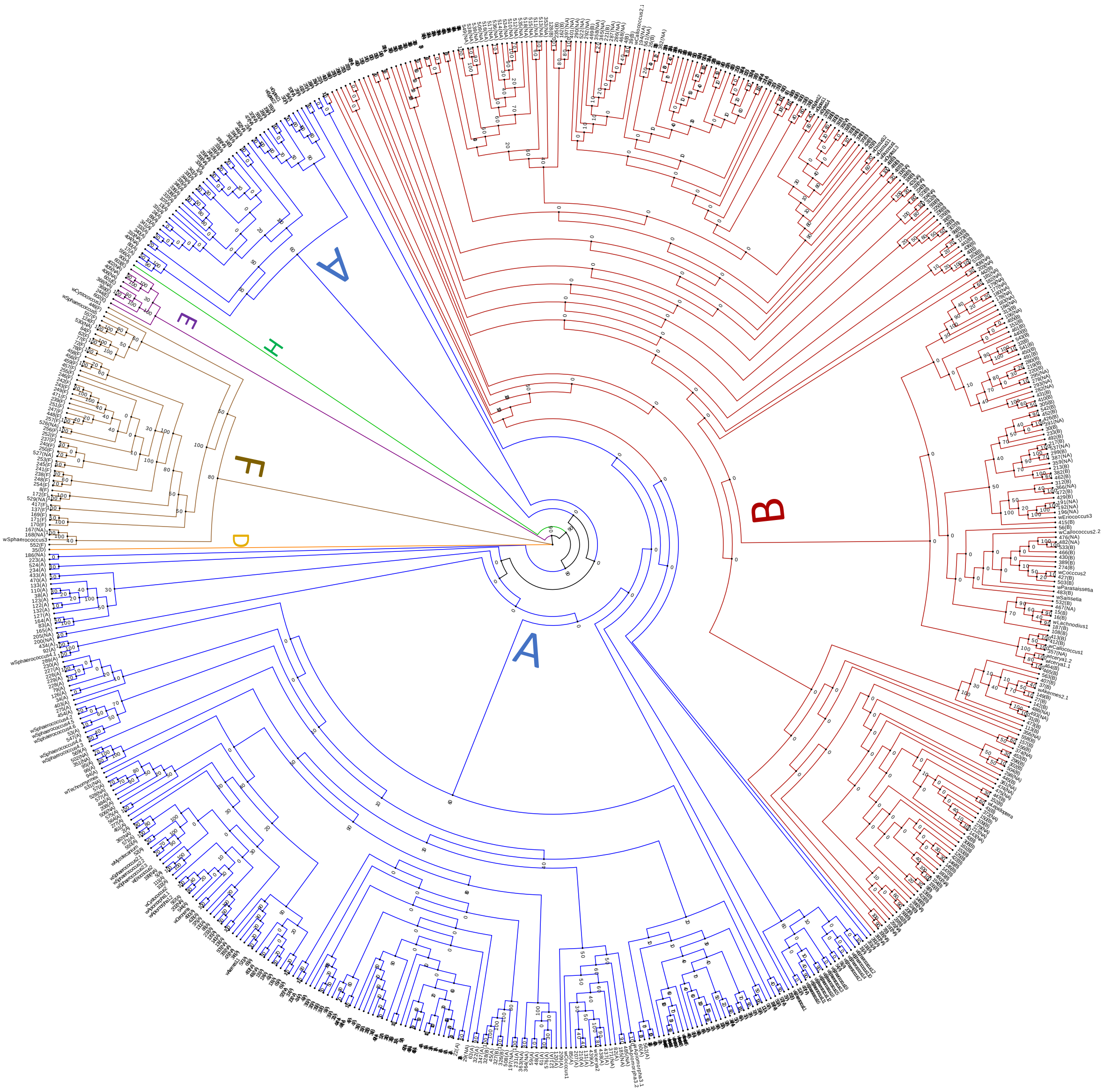


Figure S2. ML phylogenetic tree of all reported strains in *Wolbachia* MLST database plus the current study strains, categorized and coloured by supergroups. Strains that were extracted from the MLST database were assigned by their ST number following their reported supergroup in the bracket.

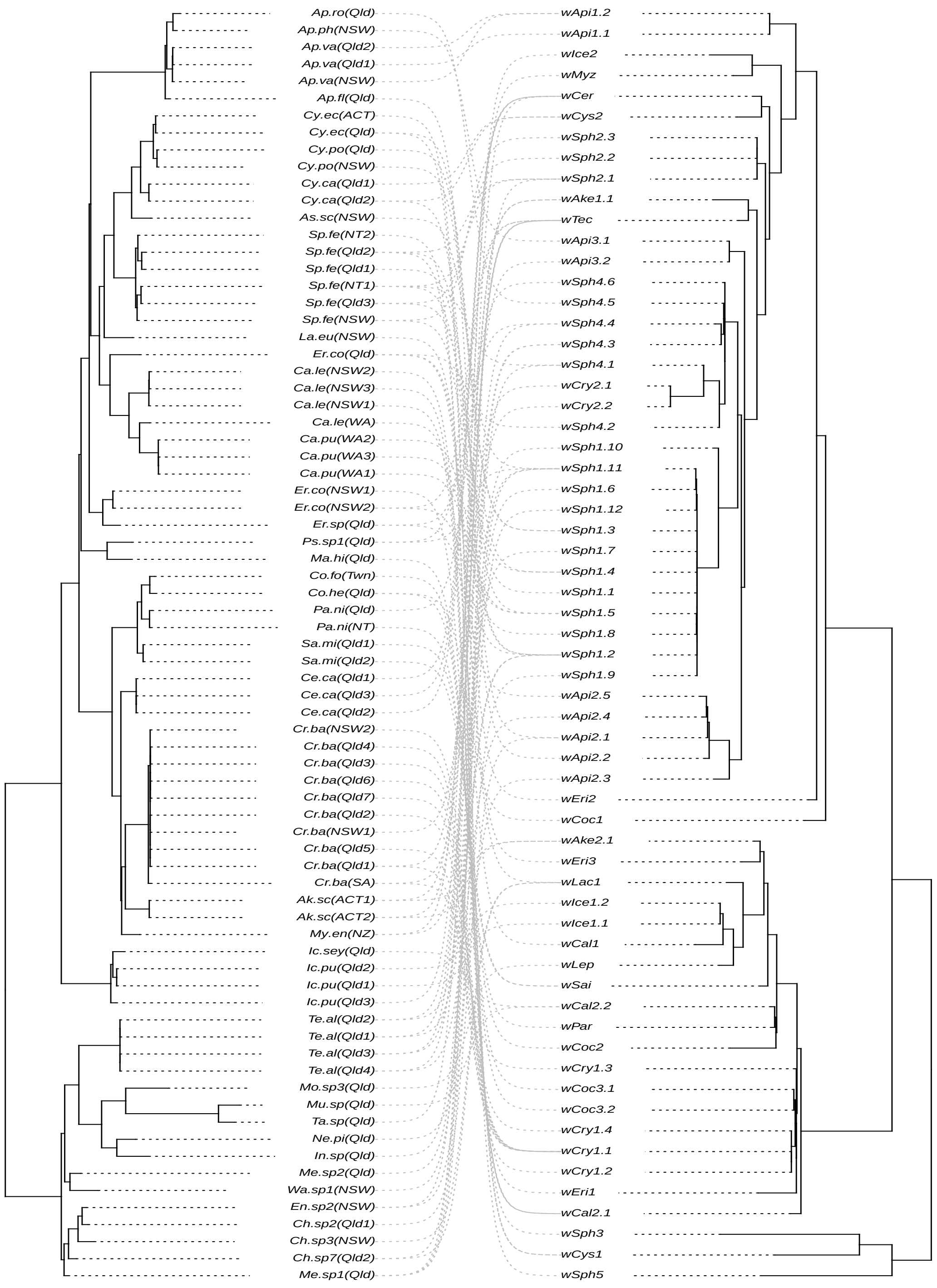


Figure S3. Phylogenetic tree of *Wolbachia* strains (on the right) and their host (scale insects and their associate species on the left). Dotted lines are indicators of *Wolbachia* infection.

#### References


Baldo L., Hotopp J.C.D., Jolley K.A., Bordenstein S.R., Biber S.A., Choudhury R.R., Hayashi C., Maiden M.C., Tettelin H. & Werren J.H. 2006. Multilocus sequence typing system for the endosymbiont Wolbachia pipientis. Appl. Environ. Microbiol. **72**: 7098–7110.

Bolger A.M., Lohse M. & Usadel B. 2014. Trimmomatic: a flexible trimmer for Illumina sequence data. Bioinformatics. **30**: 2114–2120.

Brown J., Pirrung M. & McCue L.A. 2017. FQC Dashboard: integrates FastQC results into a web-based, interactive, and extensible FASTQ quality control tool. Bioinformatics. **33**: 3137–3139.

Bushnell B., Rood J. & Singer E. 2017. BBMerge – Accurate paired shotgun read merging via overlap. PLOS ONE. **12**: e0185056.

Callahan B.J., McMurdie P.J., Rosen M.J., Han A.W., Johnson A.J.A. & Holmes S.P. 2016. DADA2: High-resolution sample inference from Illumina amplicon data. Nat. Methods. **13**: 581–583.

von Dohlen C.D. & Moran N.A. 1995. Molecular phylogeny of the homoptera: a paraphyletic taxon. J. Mol. Evol. **41**: 211–223.

Gimeno C., Belshaw R. & Quicke D.L.J. 1997. Phylogenetic relationships of the Alysiinae/Opiinae (Hymenoptera: Braconidae) and the utility of cytochrome b, 16S and 28S D2 rRNA. Insect Mol. Biol. **6**: 273–284.

Krehenwinkel H., Kennedy S.R., Rueda A., Lam A. & Gillespie R.G. 2018. Scaling up DNA barcoding – Primer sets for simple and cost efficient arthropod systematics by multiplex PCR and Illumina amplicon sequencing. Methods Ecol. Evol. **9**: 2181–2193.

Sanaei E., Lin Y.-P., Cook L.G. & Engelstädter J. 2021. Wolbachia in scale insects: a distinct pattern of infection frequencies and potential transfer routes via ant associates. 2021.08.23.457441.
