## Supplementary material for "Host phylogeny and ecological associations best explain *Wolbachia* host shifts in scale insects": File S3: File_S3.R

```
# R Scripts for Sanaei et al. 2021
# Manuscript title: Wolbachia in scale insects: genetic diversity, multiple infections, and host shifts

############################################################################################################################################################
############################################################################################################################################################
##############################################                Strain determination pipeline based on dada2            ######################################
############################################################################################################################################################
############################################################################################################################################################

# Loading libraries
if (!requireNamespace("BiocManager", quietly = TRUE))
  install.packages("BiocManager")

BiocManager::install("dada2")

path <- getwd()
list.files(path)

# Forward and reverse fastq filenames have format: SAMPLENAME_R1_001.fastq and SAMPLENAME_R2_001.fastq
fnFs <- sort(list.files(path, pattern="_trimmed_1.fastq", full.names = TRUE))
fnRs <- sort(list.files(path, pattern="_trimmed_2.fastq", full.names = TRUE))

# Extract sample names, assuming filenames have format: SAMPLENAME_XXX.fastq
# cannot have .gz file together with non-zipped file, will be imported twice here
sample.names <- sapply(strsplit(basename(fnFs), "_"), `[`, 1)
plotQualityProfile(fnFs[1:2])
plotQualityProfile(fnRs[1:2])

filtFs <- file.path(path, "filtered", paste0(sample.names, "_F_filt.fastq.gz"))
filtRs <- file.path(path, "filtered", paste0(sample.names, "_R_filt.fastq.gz"))

out <- filterAndTrim(fnFs, filtFs, fnRs, filtRs,
                     maxN=0, maxEE=c(2,2), truncQ=2, rm.phix=TRUE,
                     compress=TRUE, multithread=FALSE) # On Windows set multithread=FALSE
head(out)

errF <- learnErrors(filtFs, multithread=TRUE)
errR <- learnErrors(filtRs, multithread=TRUE)

plotErrors(errF, nominalQ=TRUE)

derepFs <- derepFastq(filtFs, verbose=TRUE)
derepRs <- derepFastq(filtRs, verbose=TRUE)
# Name the derep-class objects by the sample names
names(derepFs) <- sample.names
names(derepRs) <- sample.names

dadaFs <- dada(derepFs, err=errF, multithread=TRUE)
dadaRs <- dada(derepRs, err=errR, multithread=TRUE)
dadaFs[[1]]
# merge pairs
mergers <- mergePairs(dadaFs, derepFs, dadaRs, derepRs, verbose=TRUE)
head(mergers[[1]])

# export sequences
mergers$sequence
write.table(mergers$sequence, file="dada2_sequences.fna")

write.table(seqtab, file="dada2_sequences_counts.txt")

seqtab <- makeSequenceTable(mergers)
dim(seqtab)

# Inspect distribution of sequence lengths
table(nchar(getSequences(seqtab)))

# remove chimera
seqtab.nochim <- removeBimeraDenovo(seqtab, method="consensus", multithread=TRUE, verbose=TRUE)
sum(seqtab.nochim)/sum(seqtab)

write.table(seqtab.nochim, file="dada2_sequences_counts.txt")

# Track reads through the pipeline
getN <- function(x) sum(getUniques(x))
track <- cbind(out, sapply(dadaFs, getN), sapply(dadaRs, getN), sapply(mergers, getN), rowSums(seqtab.nochim))
# If processing a single sample, remove the sapply calls: e.g. replace sapply(dadaFs, getN) with getN(dadaFs)
colnames(track) <- c("input", "filtered", "denoisedF", "denoisedR", "merged", "nonchim")
rownames(track) <- sample.names
head(track)

## The OTUs table will export to Excel for further annotation and manual strain determination as explained in File S1. 

############################################################################################################################################################
############################################################################################################################################################
##############################################      Generating the sequence lists as an input for the phylogeny       ######################################
############################################################################################################################################################
############################################################################################################################################################
#### Loading libraries 
library(dplyr)
library(tidyverse)
library(openxlsx)

#### Data input
raw_otu <- read.xlsx("./File_S2.xlsx", sheet = "otu_table")
otu <- select(raw_otu, OTU, gene, seq)

raw_strain <- read.xlsx("./File_S2.xlsx", sheet = "Strains_congruency_test")

raw_host <- read.xlsx("./File_S2.xlsx", sheet = "Hosts_congruency_test")

#Make a subset of table including only the code of the specific gene 
#change the "Gene (eg: 18S)"  to "OTU"

OTU_18S_population <- select(raw_host, Abbr, "18S") %>% rename( OTU = "18S")
OTU_28S_population <- select(raw_host, Abbr, "28S") %>% rename( OTU = "28S")
OTU_COI_population <- select(raw_host, Abbr, "COI") %>% rename( OTU = "COI")
OTU_16S_strain <- select(raw_strain, Abbr, "16S") %>% rename( OTU =  "16S")
OTU_coxA_strain <- select(raw_strain, Abbr, "coxA") %>% rename( OTU =  "coxA")
OTU_gatB_strain <- select(raw_strain, Abbr, "gatB") %>% rename( OTU = "gatB")
OTU_fbpA_strain <- select(raw_strain, Abbr, "fbpA") %>% rename( OTU = "fbpA")
OTU_ftsZ_strain <- select(raw_strain, Abbr, "ftsZ") %>% rename( OTU = "ftsZ")
OTU_hcpA_strain <- select(raw_strain, Abbr, "hcpA") %>% rename( OTU = "hcpA")
OTU_wsp_strain <- select(raw_strain, Abbr, "wsp") %>% rename( OTU = "wsp")
### generate a new table including the gene per sample 
otu$seq <- as.character(otu$seq)

Final.18S <- merge(OTU_18S_population, otu, by = "OTU", all.x = T) %>% select(Abbr, gene, seq)
Final.28S <- merge(OTU_28S_population, otu, by = "OTU", all.x = T) %>% select(Abbr, gene, seq)
Final.COI <- merge(OTU_COI_population, otu, by = "OTU", all.x = T) %>% select(Abbr, gene, seq)
Final.16S <- merge(OTU_16S_strain, otu, by = "OTU", all.x = T) %>% select(Abbr, gene, seq)
Final.coxA <- merge(OTU_coxA_strain, otu, by = "OTU", all.x = T) %>% select(Abbr, gene, seq)
Final.gatB <- merge(OTU_gatB_strain, otu, by = "OTU", all.x = T) %>% select(Abbr, gene, seq)
Final.fbpA <- merge(OTU_fbpA_strain, otu, by = "OTU", all.x = T) %>% select(Abbr, gene, seq)
Final.ftsZ <- merge(OTU_ftsZ_strain, otu, by = "OTU", all.x = T) %>% select(Abbr, gene, seq)
Final.hcpA <- merge(OTU_hcpA_strain, otu, by = "OTU", all.x = T) %>% select(Abbr, gene, seq)
Final.wsp <- merge(OTU_wsp_strain, otu, by = "OTU", all.x = T) %>% select(Abbr, gene, seq)

### Writing files

write.csv(Final.18S,"./results/18S(new).csv", row.names=FALSE)
write.csv(Final.28S,"./results/28S(new).csv", row.names=FALSE)
write.csv(Final.COI,"./results/COI(new).csv", row.names=FALSE)
write.csv(Final.16S,"./results/16S.csv", row.names=FALSE)
write.csv(Final.hcpA,"./results/hcpA.csv", row.names=FALSE)
write.csv(Final.fbpA,"./results/fbpA.csv", row.names=FALSE)
write.csv(Final.gatB,"./results/gatB.csv", row.names=FALSE)
write.csv(Final.coxA,"./results/coxA.csv", row.names=FALSE)
write.csv(Final.ftsZ,"./results/ftsZ.csv", row.names=FALSE)
write.csv(Final.wsp,"./results/wsp.csv", row.names=FALSE)

### This files Will be aligned and re-formatted into fasta in Geneious
### Phylogenetic tree will be constructed as it is explained in the manuscript

############################################################################################################################################################
############################################################################################################################################################
##############################################     Congruence test between scale insect and Wolbachia and plotting        ###################################
############################################################################################################################################################
############################################################################################################################################################
# The origin scripts of the PACo is availabe at https://www.uv.es/cophylpaco/index.html
#### Loading libraries 
library(reshape2)
library(qdapTools)
library(openxlsx)
library(dplyr)
library(tidyr)
library(tibble)
library (ape)
library(vegan)
library(Matrix)
library(data.table)
library(phytools)
library(stringi)
library(ggtree)
library(randomcoloR)
if (!requireNamespace("BiocManager", quietly = TRUE))
  install.packages("BiocManager")

### 1. PACo FUNCTION: adjustment prior to Procrustes analysis
PACo <- function (H.dist, P.dist, HP.bin)
{ 
  HP.bin <- which(HP.bin > 0, arr.in=TRUE)
  H.PCo <- pcoa(H.dist, correction="cailliez") #Performs PCo of Host distances 
  P.PCo <- pcoa(P.dist, correction="cailliez") #Performs PCo of Parasite distances
  if (is.null(H.PCo$vectors.cor)==TRUE) H.PCo <- H.PCo$vectors else
    H.PCo <- H.PCo$vectors.cor      # returns corrected pcoord 
  if (is.null(P.PCo$vectors.cor)==TRUE) P.PCo <- P.PCo$vectors else
    P.PCo <- P.PCo$vectors.cor
  H.PCo <- H.PCo[HP.bin[,1],]  #adjust Host PCo vectors 
  P.PCo <- P.PCo[HP.bin[,2],]  #adjust Parasite PCo vectors
  list (H.PCo = H.PCo, P.PCo = P.PCo)
}
### 2. DATA INPUT
#2.1.1 Phylogenetic trees:
TreeH <- read.tree("./Host_congruencyTEST.newick")
TreeP <- read.tree("./Wolbachia_congruencyTEST.newick") 

##############Compute patristic distances:
host.D <- cophenetic (TreeH)
para.D <- cophenetic (TreeP)

#2.2 ## Read HP: host-parasite association matrix

HPconnection <- read.xlsx("./File_S2.xlsx", sheet = "network_congruency_test") 
### Plot cophylo
abj1 <- cophylo(TreeH,TreeP,rotate=FALSE,assoc=HPconnection)

palette<-distinctColorPalette(nrow(HPconnection))
palette<-make.transparent(palette,0.5)

plot(abj1,link.type="curved",link.lwd=2, link.col=palette,
     link.lty="solid",lwd= 1.5,
     link.col=make.transparent("grey",1), pts=FALSE, fsize=0.8, part=0.35)

#### Convert HPconnection it to 0,1 matrix 
HP <- as.matrix(
  HPconnection %>% 
    pivot_wider(host,
                names_from = para, 
                values_from = para, 
                values_fn = list(para = length), 
                values_fill = list(para = 0))
)
####  Host names have to be moved to column names and the matrix should be numeric
rownames(HP) <- HP[, 1] # move taxa names to rownames
HP <- HP[, -1] # delete column with taxanames

class(HP) <- "numeric"
storage.mode(HP) <- "numeric"

#Sort host and parasite taxa in distance matrices to match the HP matrix:
##########################################################

host.D <- host.D[rownames(HP), rownames(HP)]
para.D <- para.D[colnames(HP), colnames(HP)]

###### Parafit test (by using two correction options)
res_cailliez<- parafit(host.D, para.D, HP, nperm = 100000, test.links = FALSE, correction="cailliez")
res_lingoes<- parafit(host.D, para.D, HP, nperm = 100000, test.links = FALSE, correction="lingoes")

### 3. APPLY PACo FUNCTION  
PACo.fit <- PACo(host.D, para.D, HP)
HP.proc <- procrustes(PACo.fit$H.PCo, PACo.fit$P.PCo) #Procrustes Ordination 
NLinks = sum(HP) #Number of H-P links; needed for further computations
#
#3.1 Plot of host and parasite ordination:
HostX <- HP.proc$X #host ordination matrix
ParY <- HP.proc$Yrot #parasite ordination matrix, scaled and rotated to fit HostX
#Plotting host and parasite ordinations
plot(HostX, asp=1, pch=46) 
points(ParY, pch=1)
arrows(ParY[,1], ParY[,2], HostX[,1], HostX[,2], length=0.12, angle=15, xpd=FALSE)
HostX <- unique(HP.proc$X) 
ParY <- unique(HP.proc$Yrot) #unique() removes duplicated points - convenient for labeling of points below
identify(ParY[,1], ParY[,2], rownames(ParY), offset=0.3, xpd=FALSE, cex=0.8) #interactive labeling
identify(HostX[,1], HostX[,2], rownames(HostX),offset=0.3, xpd=TRUE, cex= 0.8)
#
#3.2 Goodness-of-fit-test
m2.obs <- HP.proc$ss #observed sum of squares
N.perm = 100000 #set number of permutations for testing
P.value = 0
set.seed(5)

for (n in c(1:N.perm))
{ 
  if (NLinks <= nrow(HP) | NLinks <= ncol(HP)) 	#control statement to avoid all parasites being associated to a single host 
  {	flag2 <- TRUE 
  while (flag2 == TRUE)	{ 
    HP.perm <- t(apply(HP,1,sample))
    if(any(colSums(HP.perm) == NLinks)) flag2 <- TRUE else flag2 <- FALSE
  }  
  } else { HP.perm <- t(apply(HP,1,sample))} #permutes each HP row independently
  PACo.perm <- PACo(host.D, para.D, HP.perm)
  m2.perm <- procrustes(PACo.perm$H.PCo, PACo.perm$P.PCo)$ss #randomized sum of squares
  #write (m2.perm, file = "D:/m2_perm.txt", sep ="\t", append =TRUE) #option to save m2 from each permutation
  if (m2.perm <= m2.obs)
  {P.value = P.value + 1} 
}
P.value <- P.value/N.perm
cat(" The observed m2 is ", m2.obs, "\n", "P-value = ", P.value, " based on ", N.perm," permutations.")
#
#3.3 Contribution of individual links
HP.ones <- which(HP > 0, arr.in=TRUE)
SQres.jackn <- matrix(rep(NA, NLinks**2), NLinks)# empty matrix of jackknifed squared residuals
colnames (SQres.jackn) <- paste(rownames(HP.proc$X),rownames(HP.proc$Yrot), sep="-") #colnames identify the H-P link
t.critical = qt(0.975,NLinks-1) #Needed to compute 95% confidence intervals.
for(i in c(1:NLinks)) #PACo setting the ith link = 0
{HP.ind <- HP
HP.ind[HP.ones[i,1],HP.ones[i,2]]=0
PACo.ind <- PACo(host.D, para.D, HP.ind)
Proc.ind <- procrustes(PACo.ind$H.PCo, PACo.ind$P.PCo) 
res.Proc.ind <- c(residuals(Proc.ind))
res.Proc.ind <- append (res.Proc.ind, NA, after= i-1)
SQres.jackn [i, ] <- res.Proc.ind	#Append residuals to matrix of jackknifed squared residuals
} 
SQres.jackn <- SQres.jackn**2 #Jackknifed residuals are squared
SQres <- (residuals (HP.proc))**2 # Vector of original square residuals
#jackknife calculations:
SQres.jackn <- SQres.jackn*(-(NLinks-1))
SQres <- SQres*NLinks
SQres.jackn <- t(apply(SQres.jackn, 1, "+", SQres)) #apply jackknife function to matrix
phi.mean <- apply(SQres.jackn, 2, mean, na.rm = TRUE) #mean jackknife estimate per link
phi.UCI <- apply(SQres.jackn, 2, sd, na.rm = TRUE) #standard deviation of estimates
phi.UCI <- phi.mean + t.critical * phi.UCI/sqrt(NLinks) #upper 95% confidence interval
#barplot of squared jackknifed residuals
par(mar = c(5,2,0,0))
pat.bar <- barplot(phi.mean, names.arg = " ", space = 0, col="white", ylab= "Squared residuals", ylim=c(0, max(phi.UCI)), cex.lab=1.2)
text(pat.bar, par("usr")[3] - 0.001, srt = 1000, adj = 0, labels = colnames(SQres.jackn), xpd = TRUE, font = 2, cex=0.6)
arrows(pat.bar, phi.mean, pat.bar, phi.UCI, length= 0.1, angle=90)
abline(a=median(phi.mean), b=0, lty=2, xpd=FALSE) #draws a line across the median residual value
#


############################################################################################################################################################
############################################################################################################################################################
##################################################### Generating the interactive Tanglegram  Matrix ########################################################
############################################################################################################################################################
############################################################################################################################################################
library(r2d3)
library(bipartite )
library(purrr) 
library(dplyr) 
library(tidyr) 
library(stringr)
library(tibble)
library(RColorBrewer)
library(bipartiteD3)

## Re-creat the function due to some error in the package

testdata <- data.frame(higher = c("bee1","bee1","bee1","bee2","bee1","bee3"), 
                       lower = c("plant1","plant2","plant1","plant2","plant3","plant4"), 
                       webID = c("meadow","meadow","meadow","meadow","meadow","meadow"), freq=c(5,9,1,2,3,7))
bipartite::frame2webs(testdata)-> SmallTestWeb

Used3v5<- function(data,
                   filename='bipartiteD3Script',
                   PrimaryLab = 'Primary',
                   SecondaryLab='Secondary',
                   SiteNames=NULL,
                   colouroption=c('monochrome', 'brewer', 'manual')[1],
                   HighlightLab = 'Unlinked',
                   HighlightCol = '#3366CC',
                   monoChromeCol = 'rgb(56,43,61)',
                   ColourBy = c(1, 2)[2],
                   BrewerPalette = 'Accent',
                   NamedColourVector,
                   MainFigSize = NULL,
                   SortPrimary=NULL,
                   SortSecondary=NULL,
                   mp = c(1,1),
                   MinWidth=10,
                   Pad=1,
                   IndivFigSize= c(200, 400),
                   BarSize = 35,
                   Orientation = c('vertical', 'horizontal')[1],
                   EdgeMode = c('straight','smooth' )[2],
                   BoxLabPos = NULL,
                   IncludePerc = TRUE,
                   PercentageDecimals =0,
                   PercPos = NULL,
                   CSS_Output_Supress = FALSE,
                   PRINT=FALSE){
  df<-NULL
  
  if(tibble::is_tibble(data)){
    data<- as.data.frame(data)
  }
  
  if(is.data.frame(data)){
    df<- data
  }else{
    if(is.list(data)){
      df<-List2DF(data, PrimaryLab,SecondaryLab,SiteNames)
    }else{
      dimensions<-length(dim(data))
      if(dimensions ==3){
        df<-Array2DF(data, PrimaryLab,SecondaryLab,SiteNames)
      }
      if(dimensions==2){
        df<-Matrix2DF(data,PrimaryLab,SecondaryLab, SiteNames)
      }
    }
  }
  if(is.null(df)){#
    stop('invalid data input. Valid forms are data frame, bipartite style matrix, list or array')
  }
  
  if((mp[2]*mp[1])> (ncol(df)-2)){
    warning('Making too many facets. Are you sure mp is set ok?')
  }
  if((mp[2]*mp[1])< (ncol(df)-2)){
    warning('Making too few facets. Guessing you want 1 row')
    mp[2]<-ncol(df)-2
  }
  
  if(is.null( MainFigSize)){
    MainFigSize<-c(mp[2]*700, mp[1]*700) # Having a reasonable guess at a default overall figure size
  }
  
  BP_JS_Writer(df = df,filename = filename,
               colouroption = colouroption,
               HighlightLab = HighlightLab,
               HighlightCol = HighlightCol,
               monoChromeCol = monoChromeCol,
               ColourBy = ColourBy,
               BrewerPalette = BrewerPalette,
               NamedColourVector = NamedColourVector,
               MainFigSize = MainFigSize ,
               SortPrimary = SortPrimary,
               SortSecondary = SortSecondary,
               mp = mp,
               MinWidth = MinWidth,
               Pad = Pad,
               IndivFigSize = IndivFigSize,
               BarSize = BarSize ,
               Orientation = Orientation,
               EdgeMode = EdgeMode,
               AxisLabels= c(PrimaryLab, SecondaryLab),
               FigureLabel = SiteNames,
               BoxLabPos = BoxLabPos,
               IncludePerc = IncludePerc ,
               PercentageDecimals = PercentageDecimals,
               PercPos = PercPos,
               CSS_Output_Supress = CSS_Output_Supress ,
               PRINT =  PRINT)
  
  LoadVisJS()
  
  # only line changed here to add d3_version = "5"
  r2d3::r2d3(data = NA, d3_version="5", script = paste0(filename,".js"),
             height= MainFigSize[2], width= MainFigSize[1] ,
             dependencies ="vizjs.js")
}

Used3v5(SmallTestWeb)
######################################################## Load the matrix

networks <- read.xlsx("./File_S2.xlsx", sheet = "networks_All_samples") %>% filter(webID == "Infection") %>% select(higher, lower)

# Aggregate to species level and name the columns

webmatrix <- cbind(aggregate(networks["lower"], by = networks[c("lower","higher")], FUN = length))
colnames(webmatrix)[1:3] <- c("higher", "lower", "freq")

#Creat the WebID column
webmatrix <- mutate(webmatrix, webID="Infection") 
#make the matrix by frame2webs
webmatrix_ready <- frame2webs(webmatrix, varnames = c("lower", "higher", "webID", "freq"), type.out = 
                                "list", emptylist = TRUE)
#Simple plot

bipartite_D3(webmatrix_ready)

#Adjusting Sizes to fit a complex 3D plot:
nb.cols <- 70
mycolors <- colorRampPalette(brewer.pal(8, "Set2"))(nb.cols)
mycolors <- as.character(mycolors)
mycolors <- data.frame(mycolors)
All_Samples_3DGraph <- bipartite_D3(webmatrix_ready, PrimaryLab = 'Host',
                       SecondaryLab = 'Wolbachia strain group',
                       SiteNames = 'Infection', colouroption = 'brewer',
                       MainFigSize = c(800, 1200), 
                       IndivFigSize = c(300, 900),
                       BoxLabPos = c(7, 10),
                       PercPos = c(200,200),
                       BarSize = 20,
                       MinWidth = 5,
                       Pad=5,
                       PercentageDecimals = 1,
                       filename = 'demo1')
# This plot can easily saved by "Export" function in R-Studio
############################################################################################################################################################
############################################################################################################################################################
#################################### Developing GAMM model to assess host phylogeny and geography in host-shifting #########################################
############################################################################################################################################################
############################################################################################################################################################
#Loading liberaries

library(devtools)
devtools::install_github("gfalbery/ggregplot")
install.packages("INLA",repos=c(getOption("repos"),INLA="https://inla.r-inla-download.org/R/stable"), dep=TRUE)
#Otherwise
library(tidyverse); library(ggregplot); library(igraph); library(magrittr); library(cowplot)
library(INLA); library(ggregplot); library(tidyverse); library(GGally); library(patchwork)
library(cowplot); library(gganimate); library(colorspace); library(RColorBrewer); library(MCMCglmm)
library(ggrepel);library(igraph); library(adehabitatHR); library(rgeos); library(readxl)

# theme prepration
theme_set(theme_cowplot() + 
            theme(strip.background = element_rect(fill = "white")))

AlberPalettes <- c("YlGnBu","Reds","BuPu", "PiYG")
AlberColours <- sapply(AlberPalettes, function(a) RColorBrewer::brewer.pal(5, a)[4])

AlberColours[length(AlberColours)+1:2] <- 
  RColorBrewer::brewer.pal(11, AlberPalettes[[4]])[c(2,10)]

AlberColours <- append(AlberColours, list(Pink = "#FD6396", Blue = "#3C78D8")) %>% unlist

# Loading data

Networks <- read.csv("./samples.csv")

Networks %<>% 
  rename_all(CamelConvert) %>% 
  mutate_at(c("Species"), ~.x %>% str_trim %>% str_replace_all(" ", "_"))  %>% 
  mutate_at(c("Higher"), ~.x %>% 
              str_trim %>% 
              str_replace(" ", "_") %>% 
              str_remove_all(" ")) 

Networks %>% 
  # filter(WebID == "Infection", Type == "Scaleinsect") %>% 
  rownames_to_column("ID") %>% 
  dplyr::select(Higher, Lower) %>% droplevels %>% 
  table() -> AMTable # %>% c %>% table()

AMTable %>% graph_from_incidence_matrix(directed = F) -> Graph

Graph %>% bipartite.projection() %>% extract2("proj1") %>% 
  get.adjacency(attr = "weight") %>% 
  as.matrix() ->
  
  AssMat

AssMat %>% reshape2::melt() %>% 
  rename(Wolbachia = value) ->
  Pairs

Pairs %<>% mutate_at("Wolbachia", ~as.numeric(.x>0))

# Adding distances ####

Networks %>% 
  dplyr::select(Higher, Lat, Long) %>% 
  unique %>% 
  dplyr::select(Lat, Long) %>% 
  dist %>% as.matrix -> DistMatrix

Networks %>% 
  dplyr::select(Higher, Lat, Long) %>% 
  unique %>% 
  pull(Higher) %>% list(., .) ->
  dimnames(DistMatrix)

DistMatrix %>% reshape2::melt() %>% 
  rename(Distance = value) %>% 
  left_join(Pairs, .) -> FullDF

# Importing phylogeny ####

InitialTree <- read.csv("./host_distance.csv")

InitialTree %>% pull(1) %>% 
  # str_replace_all(" ", "_") %>% 
  str_trim %>% 
  str_replace(" ", "_") %>% 
  str_remove_all(" ") %>% 
  str_replace_all(c("BotanicGardens" = "BotGdns",
                    "Qld[)]1" = "Qld)")) -> 
  TreeNames

InitialTree %<>% dplyr::select(-1)

rownames(InitialTree) <- colnames(InitialTree) <- TreeNames

HostTree <- InitialTree

HostTree %>% as.matrix %>% reshape2::melt() %>% 
  rename(Phylo = value) %>% 
  left_join(FullDF, ., by = c("Var1", "Var2")) -> FullDF2

# Importing space ####

SampleLocations <- read_xlsx("./coordinations.xlsx")

SampleLocations %<>% 
  mutate_at("Species", ~.x %>% str_trim %>% str_replace_all(" ", "_")) %>% 
  mutate_at("Location", ~.x %>% str_trim %>% str_replace_all(" ", "")) %>% 
  rename_all(CamelConvert) %>% 
  mutate(Higher = paste0(Species, "(", Location, ")"))

SampleLocations %>% 
  group_by(Species) %>% 
  summarise_at(c("Latitude", "Longitude"), mean) ->
  MeanLocations

MeanLocations %>% dplyr::select(Latitude, Longitude) %>% dist %>% as.matrix -> DistMatrix

dimnames(DistMatrix) <- list(MeanLocations$Species, 
                             MeanLocations$Species)

DistMatrix %>% reshape2::melt() %>% 
  rename(DistanceMean = value) %>% 
  left_join(FullDF2, .) -> FullDF3

FullDF3 %>% 
  mutate_at(c("Var1", "Var2"), ~str_split(.x, "[()]") %>% map_chr(1)) %>% 
  dplyr::select(Sp1 = Var1, Sp2 = Var2) %>% 
  bind_cols(FullDF2, .) ->
  FullDF3

# Making affiliates ####

Networks %>% 
  filter(WebID == "AssociateDNA") %>% 
  mutate_at("Lower", ~.x %>% str_split(" ") %>% map_chr(last))

Networks %>% 
  filter(WebID == "AssociateDNA") %>% 
  mutate_at("Lower", ~.x %>% str_split(" ") %>% map_chr(last))

# Trying an MCP ####

Coordinates <- read_xlsx("./coordinations.xlsx") %>% as.data.frame %>% 
  rename_all(CamelConvert) %>% 
  mutate_at(c("Longitude", "Latitude"), as.numeric)

Coordinates$Species %>% unique -> MCPSpecies 

MCPSpecies %>% 
  lapply(function(a){
    
    Coordinates %>% 
      filter(Species == a) %>% 
      dplyr::select(Longitude, Latitude) %>% na.omit %>% 
      SpatialPoints %>% mcp
    
  }) -> MCPList

names(MCPList) <- MCPSpecies

Dyads <- expand.grid(Sp1 = MCPSpecies, Sp2 = MCPSpecies)

Dyads %>% t %>% data.frame %>% 
  map(~MCPOverlap(MCPList, .x %>% unlist %>% as.character, Symmetrical = T)) %>% 
  unlist %>% 
  matrix(nrow = length(MCPSpecies)) ->
  HRO

dimnames(HRO) <- list(MCPSpecies, MCPSpecies)

HRO %>% reshape2::melt() %>% 
  mutate_at(1:2, ~.x %>% str_trim %>% str_replace_all(" ", "_")) %>% 
  rename(Sp1 = Var1, Sp2 = Var2, HRO = value) %>% 
  left_join(FullDF3, ., by = c("Sp1", "Sp2")) -> FullDF3

MCPList %>% map(~.x@polygons[[1]]@Polygons[[1]]@coords %>% as.data.frame) %>% 
  bind_rows(.id = "Species") ->
  PolygonDF

Coordinates %>% 
  mutate_at("Species", ~.x %>% str_trim %>% str_replace_all(" ", "_")) %>% 
  group_by(Species) %>% summarise_at(c("Longitude", "Latitude"), ~mean(.x, na.rm = T)) %>% 
  rename_at(2:3, ~paste0(.x, ".Mean")) ->
  MeanCoordinates

MeanCoordinates %>% 
  dplyr::select(Latitude.Mean, Longitude.Mean) %>% dist %>% as.matrix -> DistMatrix

dimnames(DistMatrix) <- list(MeanCoordinates$Species, 
                             MeanCoordinates$Species)

DistMatrix %>% reshape2::melt() %>% 
  rename(Sp1 = Var1, Sp2 = Var2, Distance.Mean = value) %>% 
  left_join(FullDF3, ., by = c("Sp1", "Sp2")) -> FullDF4

FullDF4 %>% 
  mutate_at(c("Var1", "Var2"), ~str_split(.x, "[(]") %>% map_chr(2) %>% 
              str_remove("[)]")) %>% 
  Grelect(Site1 = Var1, Site2 = Var2) %>% 
  bind_cols(FullDF4, .) -> FullDF4

FullDF4 %<>% mutate(SameSite = Site1 == Site2)

FullDF4 %>% saveRDS("Intermediate/FullDF.rds")


# 03_Species level models ####
# Scale Insects Model Addition ####
# devtools::install_github("gfalbery/ggregplot")

library(tidyverse); library(ggregplot); library(glue); library(magrittr); library(cowplot)

theme_set(theme_cowplot())

# Setting up the data frame and variables ####

FullDF <- readRDS("Intermediate/FullDF.rds")

Covar <- c("Phylo", "HRO", 
           "Distance")#, "Distance.Mean")

Resp <- "Wolbachia"
Family <- "binomial"

TestDF <- 
  FullDF %>%
  filter(!str_detect(Var1, "Taiwan"), 
         !str_detect(Var2, "Taiwan")) %>% 
  filter(!Sp1 == Sp2) %>% 
  dplyr::select(Var1, Var2, Sp1, Sp2, 
                SameSite,
                all_of(Covar), Wolbachia) %>% 
  group_by(Sp1, Sp2) %>% 
  summarise_at(c("Wolbachia", Covar), ~mean(.x, na.rm = T)) %>% 
  na.omit %>% droplevels %>% ungroup %>% data.frame

# TestDF %<>% mutate_at(Covar, ~c(scale(.x)))

TestDF %<>% mutate_at("Wolbachia", AsBinary)

# Adding the multi-matrix random effects ####

TestDF %<>% mutate_at(c("Sp1", "Sp2"), 
                      ~factor(.x, levels = sort(unique(c(TestDF$Sp1, TestDF$Sp2)))))

MZ1 <- model.matrix( ~ Sp1 - 1, data = TestDF) %>% as.matrix
MZ2 <- model.matrix( ~ Sp2 - 1, data = TestDF) %>% as.matrix

SppMatrix = MZ1 + MZ2

TestDF$Spp <- SppMatrix

PP <- list(Spp = list(rank = nlevels(TestDF$Sp1), 
                      diag(nlevels(TestDF$Sp1))))

AddCovar <- c("Phylo", "s(Phylo)",
              "HRO", "s(HRO)",
              # "Distance.Mean", "s(Distance.Mean)"#,
              "Distance", "s(Distance)"
)

ClashList <- list(
  
  AddCovar[1:2],
  
  AddCovar[c(1:2) + 2],
  
  AddCovar[c(1:4) + 4]
  
)

B1 <- 
  BAMModelAdd(Response = "Wolbachia", 
              Data = TestDF,
              Family = binomial(),
              Explanatory = c("Spp"), # %>% c("Phylo") %>% c("s(Phylo)"), 
              Add = AddCovar,
              # Add = AddCovar, #[c(1:5)], #[!str_detect(AddCovar, "s[(]")] %>% setdiff("Phylo"), 
              Select = T, PP = PP,
              Clashes = ClashList)

B1$FinalModel %>% summary

FitList <- list()

r = 1

Model <- 
  B1$FinalModel

# Getting model fits ####

SpCoefNames <- names(Model$coef)[str_detect(names(Model$coef), "Spp")] %>% 
  setdiff(names(Model$coef)[str_detect(names(Model$coef), "Spp2")])

SpCoef <- Model$coef[SpCoefNames]

# Effects ####
# 
HRORange <- seq(from = min(TestDF$HRO),
                to = max(TestDF$HRO),
                length = 101)  %>%
  c(mean(TestDF$HRO))
# 
# DistanceRange <- seq(from = min(TestDF$Distance.Mean),
#                      to = max(TestDF$Distance.Mean),
#                      length = 101)  %>% 
#   c(mean(TestDF$Distance.Mean))

PhyloRange <- seq(from = min(TestDF$Phylo),
                  to = max(TestDF$Phylo),
                  length = 101)  %>% 
  c(mean(TestDF$Phylo))

FitList <- expand.grid(
  HRO = HRORange,
  Phylo = PhyloRange#,
  # Distance.Mean = DistanceRange
  # SameSpecies = 0:1,
  # SameSite = 0:1
)

FitList$Spp <- matrix(0 , nrow = nrow(FitList), ncol = length(SpCoef))

FitPredictions  <- predict.gam(Model, 
                               newdata = FitList, 
                               se.fit = T)

FitList[,c("Wolbachia","Lower", "Upper")] <- 
  logistic(with(FitPredictions, cbind(fit, fit - se.fit, fit + se.fit)))

# Plotting the model effects ####

FitList %>% 
  filter(HRO == mean(TestDF$HRO)) %>%
  ggplot(aes(Phylo, Wolbachia)) + 
  geom_line() + geom_point(data = TestDF) + 
  geom_ribbon(alpha = 0.3, fill = AlberColours[[1]], 
              aes(ymax = Upper, ymin = Lower)) +
  labs(x = "Phylogenetic distance", y = "Sharing probability") +
  geom_line(colour = AlberColours[[1]]) +
  
  FitList %>% 
  filter(Phylo == mean(TestDF$Phylo)) %>%
  ggplot(aes(HRO, Wolbachia)) + 
  geom_line() + geom_point(data = TestDF) + 
  geom_ribbon(alpha = 0.3, fill = AlberColours[[1]], 
              aes(ymax = Upper, ymin = Lower)) +
  labs(x = "Home range overlap", y = "Sharing probability") +
  geom_line(colour = AlberColours[[1]]) + 
  
  ggsave("Figure1.jpeg", units = "mm", width = 150, height = 100, dpi = 600)
```
